# Evolution of quantitative traits: exploring the ecological, social and genetic bases of adaptive polymorphism

**DOI:** 10.1101/2025.06.23.661088

**Authors:** Laurent Lehmann, Charles Mullon

## Abstract

Adaptive polymorphisms in quantitative traits are characterised by the coexistence of distinct genetically encoded phenotypes maintained by selection. Such intraspecific genetic and phenotypic variation make a substantial contribution to biological diversity. Building on established results from invasion analysis, we analyse conditions under which adaptive polymorphism can emerge under long-term gradual evolution. We first note that, in general, fully characterising an adaptive polymorphism, including its number of morphs, trait values and frequencies, is a computationally hard problem. Nevertheless, standard invasion results reveal a unifying necessary condition for the gradual emergence of polymorphism: negative trait-dependent selection (NTDS), whereby the marginal fitness effect of increasing trait expression in a mutant depends negatively on the population-level trait expression, reflecting a negative interaction between mutant and population trait changes. We use NTDS as an organising principle to structure the pathways through which polymorphism can evolve. This is based on a decomposition of fitness that distinguishes production factors, the inputs to survival and reproduction that are endogenous to individuals, from regulating factors, the ecological, social or genetic inputs to fitness that are influenced in part by conspecifics. This decomposition allows us to clarify how NTDS arises across three broad classes of interactions: ecological, social and genetic; and to identify previously unrecognised adaptive polymorphisms in well-studied scenarios, including life-history, social preferences, and class-structured populations. More broadly, by integrating perspectives from population genetics, evolutionary ecology, adaptive dynamics, game theory and inclusive fitness theory, we aim to provide a common thread among the pathways through which selection favours intraspecific variation.

## 1 Introduction

Polymorphism–the coexistence of multiple genetically determined phenotypes within a population–is widespread in nature, from discrete colour morphs in many flowering plants and animals (e.g. Gigord et al., 2001; Sinervo and Lively, 1996), to alternative mating strategies in vertebrates (e.g. Gross, 1996; Oliveira et al., 2008), sexual dimorphism in many species (e.g. Lande, 1980; Fairbairn, 2013), and allelic diversity at metabolic loci (e.g. Gillespie, 1991; Tishkoff et al., 2001). Unlike polyphenism, where distinct phenotypes are environmentally induced by the same genotype (West-Eberhard, 2003), polymorphism necessarily involves genetic variation, maintained by the segregation of different alleles. Although non-selective processes such as mutation, recombination, and genetic drift shape genetic variation (Crow and Kimura, 1970; Lynch and Walsh, 2018), natural selection is the only evolutionary force that can maintain cohesive and clearly differentiated phenotypes over evolutionary timescales. Here, we use “adaptive polymorphism” to refer to an intraspecific polymorphism that is underlain by the segregation of different alleles whose coexistence is favoured by natural selection and by natural selection alone (thus resulting in the stable maintenance of phenotypic variation in the absence of mutation). Understanding the conditions that favour adaptive polymorphism is important for identifying the mechanisms that promote and maintain the diversity of forms and functions so characteristic of the natural world.

Different theoretical frameworks, including population genetics, evolutionary ecology, and evolutionary game theory, have proposed a variety of mechanisms that favour intraspecific adaptive polymorphism. These include temporally or spatially varying selection (e.g. Levene, 1953; Gillespie, 1991; Svardal et al., 2015; Yamamichi et al., 2023), pleiotropy and trade-offs between competitive traits due to functional constraints (e.g. de Mazancourt and Dieckmann, 2004; Ito and Sasaki, 2016), niche differentiation (e.g. Karlin and McGregor, 1972), strategic interactions producing frequency-dependent fitness effects in social contexts (e.g. Maynard Smith, 1982; McNamara and Leimar, 2020), and balancing selection via heterozygote advantage (e.g. Karlin and McGregor, 1972; Lewontin, 1974; Hartl and Clark, 2007; Sellis et al., 2011). This diversity of mechanisms can make it difficult, when analysing a specific biological scenario, to determine which intuition to apply or what general conditions, if any, govern the emergence of adaptive polymorphism.

A common feature unifying these diverse mechanisms is that some form of negative intraspecific population feedback must act to favour diversity. However, this feedback differs across theoretical frameworks. Evolutionary ecology models often invoke environmental feedbacks, mediated by density-dependent resource depletion, co-adapted parasitism, or prey-predator interactions (e.g. Geritz et al., 1998; Pásztor et al., 2016). Game-theoretic models emphasise social feedback and frequency-dependent selection, where the payoff to behavioural strategies, such as aggression, cooperation, or signalling, depends on their frequency in the population (e.g. Hofbauer and Sigmund, 1998; McNamara and Leimar, 2020). Population genetics, meanwhile, highlights genetic feedbacks, notably overdominance in diploids, where allelic fitness depends directly on allele frequencies within genotypes and thus on mating patterns (e.g. Crow and Kimura, 1970; Karlin and McGregor, 1972; Ewens, 2004).

How do these various population feedbacks influence the emergence of adaptive polymorphism when new phenotypes regularly arise by mutation? What are the relationships between these different mechanisms? We address these questions in the context of quantitative traits evolving gradually through Darwinian adaptive dynamics, i.e. through the many successive small trait modifications. We build on the established theory of analysing the emergence of polymorphism under gradual evolution (e.g. Taylor, 1989; Christiansen, 1991; Metz et al., 1996; Eshel et al., 1997; Vincent et al., 1996; Geritz et al., 1998; Ajar, 2003; Rousset, 2004; Vincent and Brown, 2005; Otto and Day, 2007; Dercole and Rinaldi, 2008; Sasaki and Dieckmann, 2011; Brännström et al., 2013; Kisdi and Geritz, 2016; Mullon et al., 2016; Avila and Mullon, 2023), adopting evolutionary invasion analysis or long-term evolution theory (“ESS theory”, see Van Cleve, 2023, for review). This provides a unifying framework to study conditions under which adaptive polymorphisms initially emerge from ecological, social, and genetic feedbacks.

Our study proceeds as follows. We begin by formalizing the concept of an adaptive polymorphism and show its useful equivalence with the concept of Nash equilibrium of a population game. This correspondence highlights that mean invasion fitness is maximized at an adaptive polymorphism in a sense to be made precise below. It is, however, computationally intractable in general to characterize this polymorphism explicitly. We therefore turn to classical invasion analysis, which identifies negative trait-dependent selection (NTDS) as a key necessary condition for the gradual emergence of polymorphism, whereby changes in mutant trait value and changes in the population trait value interact non-additively in determining mutant fitness, with higher population trait values reducing the marginal fitness gain from further increases in that trait. We next examine the population feedbacks that generate NTDS by expressing fitness in terms of production and regulating factors, which link heritable trait values to survival and reproduction through their genetic, social and ecological effects. Building on this, we analyse how NTDS arises across three main classes of biological interactions: ecological, social, and genetic; and derive the conditions under which each can promote adaptive polymorphism. Finally, we apply these conditions to classical models of quantitative trait evolution, revealing previously overlooked polymorphisms.

## 2 Evolutionary model

### 2.1 Biological assumptions

We consider a large (ideally infinite) density-dependent regulated population. We start by assuming that individuals are haploid and reproduce asexually (this seemingly strong assumption is relaxed in section 6.1, where we consider diploidy and sexual reproduction). Individuals can belong to a finite number of different classes (e.g. be of different ages or be in different physiological states) and can be distributed in groups provided there is no isolation by distance (e.g. group or patch structure according to the infinite island model of dispersal, Wright, 1931; Rousset, 2004). The population may also experience environmental fluctuations in space and time as long as the biotic and abiotic processes driving these fluctuations are stationary ergodic. We census this population at discrete demographic time steps between which individuals can survive, reproduce and interact.

Each individual expresses a quantitative or continuous trait; that is, a trait whose heritable value belongs to a subset 𝒳 ⊆ ℝ of the real numbers. This trait can influence any aspect of an individual’s biology, from its morphology to its physiology and behavior. We assume that the trait evolves gradually via the constant influx of rare genetic mutations of small effects.

### 2.2 Adaptive polymorphism

#### 2.2.1 Background concepts and invasion fitness

We are interested in describing what constitutes an adaptive polymorphism in the evolving trait. To that end, we introduce several concepts. The first is the probability distribution *p* of trait values segregating in the population at a given demographic time point (*p* is an element of Δ(𝒳), the set of probability distributions over the trait space 𝒳). For a finite number of morphs *p*(*x*) can be thought of as the frequency of the allele encoding trait value *x* ∈ 𝒳. More generally, we do not assume that *p* has a density (with respect to the Lebesgue measure) in order to be able to cover both the cases of a finite and infinite number of different segregating trait values. The probability distribution *p* is the main quantity used to characterise the state of a population in evolutionary biology, with its change in time constituting the basis of evolutionary dynamics (e.g. Wright, 1931; Hamilton, 1964; Crow and Kimura, 1970; Nagylaki, 1992; Charlesworth, 1994; Bürger, 2000, and see Bürger, 2000, chapter 4 for covering both finite and infinite number of segregating morphs). However, *p* alone is seldom sufficient to characterize evolutionary dynamics as these typically rely on several additional state variables (e.g., population size, environmental variables, genotype frequencies for diploids, Crow and Kimura, 1970; Nagylaki, 1992; Charlesworth, 1994; Bürger, 2000). All the auxiliary state variables that are necessary to close the dynamics of the distribution *p*, we collect in *ζ*. At any demographic time step, the state of the population is therefore fully described by *ω* = (*p, ζ*).

Following Ellner and Sasaki (1996), we assume that the evolutionary process in the absence of mutation is such that given any set of alleles initially segregating (i.e. given any subset of segregating morphs in 𝒳), the population state *ω* converges towards an attractor that is stationary ergodic (thus excluding the existence of multiple attractors). We denote this attractor as *µ*, i.e. *µ* is the stationary distribution of *ω* over all states the population can be in. Thus, *µ* is the joint distribution of *p* and *ζ*, and we denote by *µ*_p_ the marginal stationary ergodic distribution of *p*. Under a stationary ergodic process, an allele is either present or absent at all demographic time points. Let us then denote by 𝒫(*µ*_p_) the set of alleles (and thus morphs) present in the population under the distribution *µ*_p_.

A population is polymorphic when 𝒫(*µ*_p_) contains two or more traits (or morphs). If the population is monomorphic for *x*, then 𝒫(*µ*_p_) = {*x*}. If two trait values *x*_1_ and *x*_2_ coexist, then 𝒫(*µ*_p_) = {*x*_1_, *x*_2_}. More generally, the set 𝒫(*µ*_p_) of segregating traits can be finite, consisting of a finite number of traits (i.e. an oligomorphism *sensu* Sasaki and Dieckmann, 2011), *or infinite consisting of either a continuum of traits or an infinite number of discrete traits. A population is polymorphic when 𝒫(µ*_p_) contains two or more trait values (or morphs).

Let us now consider a mutant type *x* introduced as a single copy in a resident population characterized by *µ* and thus trait distribution *µ*_p_, where *µ* can be thought of as the environment from the point of view of the mutant. Under our assumptions of section 2.1, the dynamics of the number of mutant copies descending from the initial mutant can be modelled as a multitype branching process (e.g., Harris, 1963; Mode, 1971 for constant environments and Tanny, 1981; Vatutin, 2021 for random environments). We assume that this process satisfies an *extinction dichotomy* defined by two features. First, the process is characterized by either the event of sure extinction occurring with probability one or the event of non-sure extinction occurring with probability one (e.g. Vatutin, 2021, eq. 15). Biologically, this entails that the mutant lineage—started in any possible population state with a single mutant copy—goes extinct with probability one or has a positive probability of invading (i.e., all states with a finite number of mutants are transient). Second, the event of sure extinction occurs with probability one if and only if

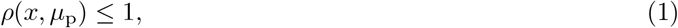

where *ρ*(*x, µ*_p_) is the long-term geometric mean of the average per-capita number of mutant copies produced per demographic time step in a population where the segregating trait values are in 𝒫(*µ*_p_). Since it determines the fate of the mutant, we refer to *ρ*(*x, µ*_p_) as the invasion fitness of mutant *x* in resident population *µ*_p_. We have written invasion fitness as depending explicitly only on *µ*_p_ rather than on the full environment *µ*. We can do this because the set 𝒫(*µ*_p_) of segregating morphs determines *µ* (on knowledge of the dynamics of *µ*), and this set is itself determined by *µ*_p_.^1^ The dependence of invasion fitness on *µ* is thus left implicit as our aim is not to compute invasion fitness explicitly at this stage, but to use it to provide a characterisation of an adaptive polymorphism.

There is a wide class of situations satisfying the extinction dichotomy. We do not detail here the sufficient conditions for this (see Tanny, 1981, Th. 9.6 and Th. 9.10 and Vatutin, 2021, Th. 1 and Th. 2), but note that these extend the usual conditions for constant environments, namely: that the mutant fitness matrix is primitive (e.g. Tuljapurkar, 1990; Caswell, 2001) and that the branching process is non-singular (i.e., each individual does not produce exactly one successful offspring; Harris, 1963, p. 38–41; see Box 3 for an example of a fitness component underlying invasion fitness, and Lehmann, 2026 for more details on the extinction dichotomy and invasion fitness under our island model assumptions). When the mutant invades, invasion fitness is equivalent to the long-term geometric growth ratio of the mutant lineage size (point 3 of Tanny, 1981, Th. 9.6). As such, fitness *ρ*(*x, µ*_p_) captures the various concepts of invasion fitness used in invasion analyses (e.g., Fisher, 1930; Eshel and Feldman, 1984; Altenberg and Feldman, 1987; Liberman, 1988; Lessard, 1990; Tuljapurkar, 1990; Metz et al., 1992; Charlesworth, 1994; Arnold et al., 1994; Rand et al., 1994; Ferrière and Gatto, 1995; Ellner and Sasaki, 1996; Metz et al., 1996; Vincent and Brown, 2005; Otto and Day, 2007; Metz, 2011; Lehmann et al., 2016; Altenberg et al., 2017; Van Cleve, 2023).

#### 2.2.2 Formalizing adaptation and biological interpretation

We now define an adaptive polymorphism as: a distribution 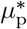 such that (i) the set of segregating morphs 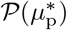 contains more than one morph; and (ii) this coalition of morphs is uninvadable and thus satisfies

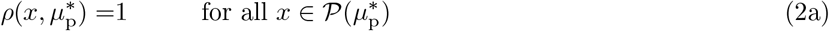

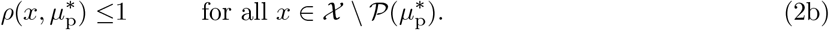

Condition (2a) entails that each morph 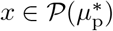 in the uninvadable coalition exactly replaces itself through survival and reproduction, and is therefore maintained in the population. Condition (2b), meanwhile, ensures that any type 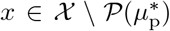 outside of the coalition goes extinct if introduced as a single copy into the population. The definition of an evolutionary stable strategy by Ellner and Sasaki (1996, p. 39) is equivalent to eq. (2), except that they impose a strong inequality in eq. (2b). The strong inequality, however, is not necessary where mutations are assumed to arrive as single copies in a large population where a branching process can be used to determine their fate (i.e., the extinction dichotomy holds).

When both conditions (2a)–(2b) hold, the polymorphism can no longer be altered through natural selection on the introduction of alternative morphs by mutation. Combining both conditions shows that for each 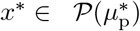, morph *x*\* solves the maximization problem 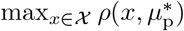 (i.e. each 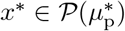 is a best response to the uninvadable coalition). Hence, each morph in the uninvadable coalition satisfies the verbal biological interpretation of an adaptation: “a phenotypic variant that results in the highest fitness among a specified set of variants in a given environment” (Reeve and Sherman, 1993, p. 9). This biological interpretation of adaptation also suggests that in an adaptive polymorphic population state, population fitness should be maximized. This turns out to be the case and to see what is meant by this, consider the case of no temporal fluctuations, in which case 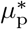 puts all probability mass on a single distribution at all demographic time step, denoted *p*\* (see Box 1 for the general case). We can then replace 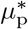 by *p*\* in eq. (2) and let 𝒫(*p*\*) denote the set of morphs present in the population under *p*\*. Then, the distribution *p*\* is an adaptive polymorphism if:

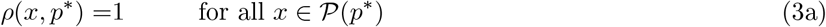

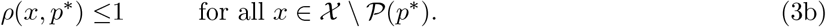

Let us now denote by

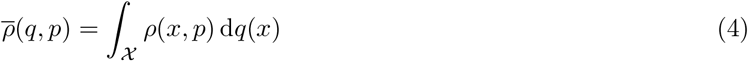

the mean invasion fitness of a mutant coalition *q* in a population where the resident coalition is *p* (here ∫_𝒳_ *g*(*x*) d*q*(*x*) denotes the expectation of *g*(*x*) over the probability measure *q* and ∫ _𝒳_ d*q*(*x*) = 1). That is, 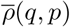 gives the mean invasion fitness over the morphs that are part of a coalition *q* in a resident population characterised by *p*. Existing results then allow us to say that *p*\* satisfies eq. (3) if and only if it satisfies

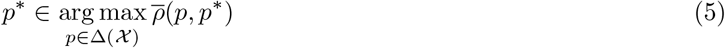

(Anderson et al., 2026, Proposition 1). In other words, *p*\* is an uninvadable coalition if and only if *p*\* maximizes mean invasion fitness at the population state *p*\* (this extends to fluctuating environments, see Box 1). No other coalition has greater mean invasion fitness than *p*\* when the population is characterised by *p*\*. Thus, in an uninvadable population state, all variation decreasing mean invasion fitness has been removed: it is as if the population state is at an optimum for trait diversity (given by *p*).

#### 2.2.3 The challenge of characterising an adaptive polymorphism

An adaptive polymorphism may consist of a finite or infinite number of morphs. Yet previous results suggest that long-term evolution should lead to a finite number of segregating morphs; namely, to an oligomorphism (*sensu* Sasaki and Dieckmann, 2011). In particular, Ellner and Sasaki (1996) show that when invasion fitness is an analytic function, the uninvadable polymorphism can only be discrete. This is in line with the finding that the equilibria of the continuum replicator dynamics in the absence of mutation consist of a finite number of coexisting morphs in panmictic populations without class structure (van Veelen and Spreij, 2009; Hingu et al., 2008). More generally, Gyllenberg and Meszéna (2005) demonstrate that the coexistence of an infinite number of morphs along a continuous range of trait values is not robustly stable, as it is sensitive to parameter changes. Hence, an adaptive polymorphism (if any) should ultimately be an oligomorphism.

Ideally, one would be able to characterize such an uninvadable oligomorphism in a quantitative way for any given biological scenario; that is, one would be able to tell the number of morphs in the coalition, the traits they express, and their frequency in the population. Moreover one would like to show that such an equilibrium is evolutionarily attainable, i.e. is convergence stable (Eshel, 1983; Eshel et al., 1997; Leimar, 2009). Working towards these general goals, however, proves especially challenging, even in the absence of temporally fluctuating selection, class structure, and under the assumption that the population state converges to a single attractor.

The challenge is revealed by the equivalence between eqs. (3) and (5), which establishes a connection between an uninvadable coalition and a Nash equilibrium of a population game (e.g. Sandholm, 2011; Anderson et al., 2026). This connection provides us with two insights. First, in the absence of temporal fluctuations, an uninvadable coalition exists (under the assumptions that invasion fitness is continuous in its arguments and 𝒳 is a closed interval, Appendix A). Second, characterizing the uninvadable coalition–computing the Nash equilibrium–is likely to be a computationally hard problem. Hence, characterising the long-term equilibrium can be intractable, even leaving aside the questions of the evolutionary convergence towards adaptive polymorphism and the complication of population heterogeneity.

To circumvent these difficulties, we next focus on one relevant segment of the evolutionary dynamics and investigate the conditions that are required for adaptive polymorphism to emerge from a population initially consisting of a single morph.

### 2.3 Monomorphic instability through invasion analysis

#### 2.3.1 Gradual emergence of polymorphism

The mathematical machinery for determining when a monomorphic population becomes polymorphic through gradual evolution is well-established (e.g. Taylor, 1989; Christiansen, 1991; Ellner and Sasaki, 1996; Metz et al., 1996; Eshel et al., 1997; Geritz et al., 1998; Rousset, 2004; Otto and Day, 2007; Dercole and Rinaldi, 2008; Sasaki and Dieckmann, 2011; Brännström et al., 2013; Kisdi and Geritz, 2016; Mullon et al., 2016). We summarise the established results needed for our analysis here. The starting point is the invasion fitness of a mutant with trait value *x* in a resident population monomorphic for trait value *y*. Since the resident population is monomorphic (i.e. 𝒫(*µ*) = {*y*}), we from now on use the notation *ρ*(*x, y*) to represent the invasion fitness of mutant *x* in a *y* population. We assume that *ρ*(*x, y*) is differentiable with respect to both of its arguments. Owing to density-dependent regulation, the invasion fitness of a single copy of any resident *y* ∈ 𝒳 must be one, *ρ*(*y, y*) = 1; that is, each gene copy on average replaces itself in a monomorphic population at its demographic attractor.

Whether a population monomorphic for *y* is evolutionarily unstable and whether such instability favours polymorphism can be examined from the following three coefficients of selection:

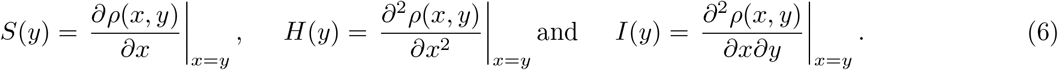

We refer to these as the selection gradient (*S*), the disruptive selection coefficient (*H*), and the selection interaction coefficient (*I*), respectively. The selection gradient *S*(*y*) indicates the direction of selection on the trait in a population at *y*. When *S*(*y*) *>* 0 (*S*(*y*) *<* 0), a mutation with small positive (negative) effect on trait value has a positive probability of invading, and if it does invade, it fixes and substitutes the resident (Eshel et al., 1997; Rousset, 2004; Cai and Geritz, 2020; Priklopil and Lehmann, 2021). Therefore, *S*(*y*) is sufficient to characterise gradual evolution under the constant but rare input of mutations with small effects: the population will converge either to a boundary point of the state space 𝒳, or to a trait value *y*\* lying in the interior of 𝒳 such that

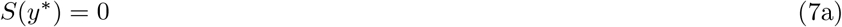

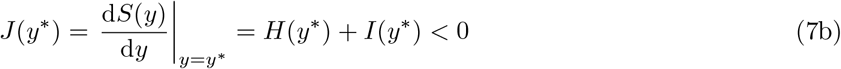

(e.g. Eshel, 1983; Taylor, 1989; Eshel et al., 1997; Leimar, 2009). A trait value *y*\* satisfying the condition in eq. (7a) is known as a singular trait value, while those satisfying also the condition in eq. (7b) are further said to be convergence stable. The coefficient *H*(*y*\*), meanwhile, indicates whether selection is stabilizing or disruptive when all individuals in a population express a singular trait value *y*\*. When *H*(*y*\*) *<* 0, any change in trait value decreases invasion fitness, so that selection ousts any mutant whose value deviates locally from *y*\*, i.e. selection is stabilizing for *y*\*. In contrast, when *H*(*y*\*) *>* 0 any trait change increases fitness so that selection favours any mutant whose value deviates locally from *y*\*, i.e. selection is disruptive. When a convergence stable trait value *y*\* is such that *H*(*y*\*) *>* 0, the population first converges to *y*\* and then gradually splits into two morphs (Eshel et al., 1997; Geritz et al., 1998). This represents the gradual emergence of polymorphism due to selection in a process often referred to as evolutionary branching (yet polymorphism is not necessarily permanent after branching, Geritz et al., 1999).

#### 2.3.2 Negative-trait dependent selection (NTDS) is necessary for polymorphism

It follows that two conditions must hold for a monomorphic population to evolve an adaptive polymorphism: *H*(*y*\*) + *I*(*y*\*) *<* 0 (for convergence stability) and *H*(*y*\*) *>* 0 (for disruptive selection). Together, these imply that

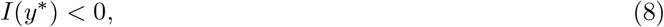

whereby the selection interaction coefficient must be negative at *y*\*. The selection interaction coefficient measures how the marginal fitness effect of changing the mutant trait value (as in *∂ρ*(*x, y*)*/∂x*) varies with the trait value *y* expressed in the resident population (recall eq. 6). In other words, it quantifies how selection depends on the resident population’s trait value; we refer to this dependence as “trait-dependent selection”. Condition (8) implies that negative trait-dependent selection (NTDS) is necessary for adaptive polymorphism to emerge by gradual evolution. Biologically, *I*(*y*\*) *<* 0 means that mutant invasion fitness increases when mutant and resident trait values vary in opposite directions.^2^ It indicates that the mutant’s best response to individuals in the population doing more of something (e.g., increase *y*), is to do less (e.g., decrease *x*), i.e. mutant and population trait values are strategic substitutes *sensu* game theory (Bulow et al., 1985) (sometimes described as an “antagonistic interaction” in biology).

The relevance of condition (8) is underscored by two further results. First, it is also the local requirement for mutual invasibility: two types that are weakly diverged around a singularity *y*\* can invade one another and thus coexist in a protected polymorphism only if *I*(*y*\*) *<* 0 (Eshel et al., 1997; Geritz et al., 1998, and see Appendix B here). That is, condition (8) is necessary for negative-frequency dependent selection near *y*\* (Maynard Smith, 1998; Heino et al., 1998). Second, once *I*(*y*\*) *<* 0 holds, it is in principle always possible to define a trade-off or add an individual cost of trait expression such that *H*(*y*\*) *>* 0 and *J*(*y*\*) *<* 0 (Kisdi, 2006, 2015; Ito and Sasaki, 2016).

While our focus here is on the conditions for instability of monomorphism, the same principles can be extended to study the instability of polymorphism. Examining invasion fitness and selection coefficients, this time in polymorphic resident populations, allows one to determine whether a given oligomorphism is convergence stable and uninvadable (e.g. Geritz et al., 1998; Ito and Dieckmann, 2007; Sasaki and Dieckmann, 2011). In such analyses, NTDS remains the key diagnostic quantity, capturing the negative population feedback on which adaptive polymorphism ultimately rests (Appendix C for details). In what follows, we use *I*(*y*\*) as a unifying measure to delineate the bases of adaptive polymorphism.

## 3 Fitness factors: production, regulation, and their interactions

To specify the negative population trait feedback embodied by *I*(*y*\*) *<* 0 and better understand the biological processes via which it can be generated, we need to be more explicit about what factors influence individual survival and reproduction–”fitness factors” for short–and thus contribute to invasion fitness. From its definition, invasion fitness *ρ*(*x, y*) can be seen as the average of the individual fitness of mutants. This average, which may be arithmetic, geometric, or a combination of the two depending on the biological situation, needs to be taken over all the relevant contexts in which the gene that codes for *x* may find itself; that is, all types of genetic (e.g. genomic background), demographic (e.g. population size), ecological (e.g. prey density), and environmental states (e.g. temperature, see examples below and Box 2). Invasion fitness may thus be influenced by an untold number of factors. Yet, these can often be usefully grouped into different categories, as we next outline.

### 3.1 Limiting, regulating, and production factors

Fitness factors that have always been considered fundamental are those limiting survival and reproduction, i.e. limiting factors (Grovers, 1997). These fall into two types. On the one hand, limiting factors may be exogenous, such as temperature or humidity. On the other hand, limiting factors can be endogenous, meaning that their availability depends on conspecifics, such as their abundance and the traits they express. Examples include available space or nutrients. These endogenous limiting factors are special cases of a broader category usually referred to as *regulating factors* (e.g. May, 1973; Armstrong and McGehee, 1980; Grovers, 1997; Pásztor et al., 2016). Regulating factors are defined here as inputs to individual survival and reproduction that depend directly or indirectly on the traits expressed by *other* individuals in the population. Regulating factors are typically thought to have negative effects on individual fitness, for instance through competition for limited resources (Grovers, 1997). However, regulating factors may also enhance fitness, as occurs when mutualistic interactions improve resource availability. In this paper, regulating factors represent the first of two broad categories of fitness factors we explicitly consider.

The second category, which we refer to as *production factors*, comprises inputs to individual survival and reproduction that depend, at least partially, on the trait *x* of its bearer. Unlike regulating factors, which depend solely on others, production factors reflect intrinsic characteristics of an individual, such as baseline fecundity in the absence of competition, germination probability, energy absorption rates, or behavioral predisposition. As these examples suggest, production factors are typically associated with improved survival and/or reproduction. However, production factors of one individual often affect the regulating factors experienced by others, creating indirect interactions among individuals. These interactions between regulating and production factors may combine in complex ways to shape overall individual reproduction and survival. To disentangle these effects and explore their evolutionary implications, we next introduce a model explicitly linking trait *x* to its invasion fitness through a structured hierarchy of regulating and production factors.

### 3.2 Invasion fitness in terms of regulating and production factors

The first production factor we consider is the phenotype, defined broadly here as any physiological, morphological, or behavioural phenotype expressed by an individual (e.g. nutrient uptake rate, coat colour, effort into alloparental care). The phenotype expressed by an individual carrying trait *x* may depend not only on *x* but also on its direct or indirect interactions with others in the population. To capture this, we write the *n*_z_-dimensional phenotype **z** of an individual as a function 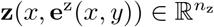 that depends on: (i) the heritable trait value *x* it carries (first argument); and (ii) a set of *r*_z_ regulating factors, which can be formalised as a function 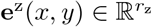 of mutant *x* and resident *y* trait values

For example, in a model of social interactions where individuals can respond to each other’s actions (e.g. Akçay and Van Cleve, 2012), the behaviour of a mutant carrier depends on the trait of its social partners (section 5.2 for examples). These interaction partners may be residents (under random interactions), or possibly other mutants (under preferential interactions among relatives or limited dispersal; section 6.2 for examples). Similarly, in diploid populations, this formulation allows for gene interactions within individuals on the phenotype, where mutant carriers are heterozygotes under random mating but may be homozygotes (carrying two copies of *x*) under inbreeding (section 6.1 for more details on diploids). More generally, **z**(*x*, **e**^z^(*x, y*)) allows us to capture biotic and abiotic regulating factors that determine the expression of quantitative traits, including environmental effects, non-additive genetic effects, gene-by-environment effects, and indirect genetic effects (Lynch and Walsh, 2018).

The second production factor we consider explicitly is termed payoff, an umbrella term for all the material and informational inputs that are acquired by the individual and that serve for its survival or reproduction (e.g. number of mates, prey caught, quality of breeding spot, calories, knowledge, Maynard Smith, 1982; Stephens and Krebs, 1987; Houston and McNamara, 1999). We conceive the *n*_*π*_-dimensional payoff of an individual as a function 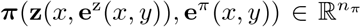 that depends on: (i) the phenotype **z**(*x*, **e**^z^(*x, y*)) it expresses (first argument); and (ii) a number *r*_*π*_ of regulating factors, given by a function 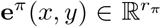 of mutant and resident trait values. Such regulating factors would include the behavioural phenotype of others that the individual interacts with, who may also be mutant carriers where interactions occur in the family or in dispersal-limited groups, or whose behaviour depends on the individual’s own trait under social responsiveness.

Finally, the translation of payoff into actual survival and reproduction may itself depend on further regulating factors that we collect in 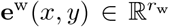. These additional *r*_w_ regulating factors include every biotic and abiotic factors outside the individual that may influence its survival and reproduction and that are affected by conspecifics (holding the phenotype and payoff of the individual fixed). Examples include population density, the payoff of competitors, or prey density. In a well-mixed population, such regulating factors would typically depend on the resident trait *y*, but under limited dispersal, they may also depend on the mutant *x* (e.g. Rousset and Ronce, 2004; Mullon and Lehmann, 2018). Putting it all together, we obtain that the invasion fitness of the mutant *x* in a resident population characterised by trait *y* can be written as

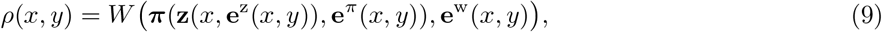

where the function *W* has as its first argument the payoff ***π*** and second argument the regulating factors **e**^w^(*x, y*) (see Fig. 1 for an illustration).

**Figure 1.**
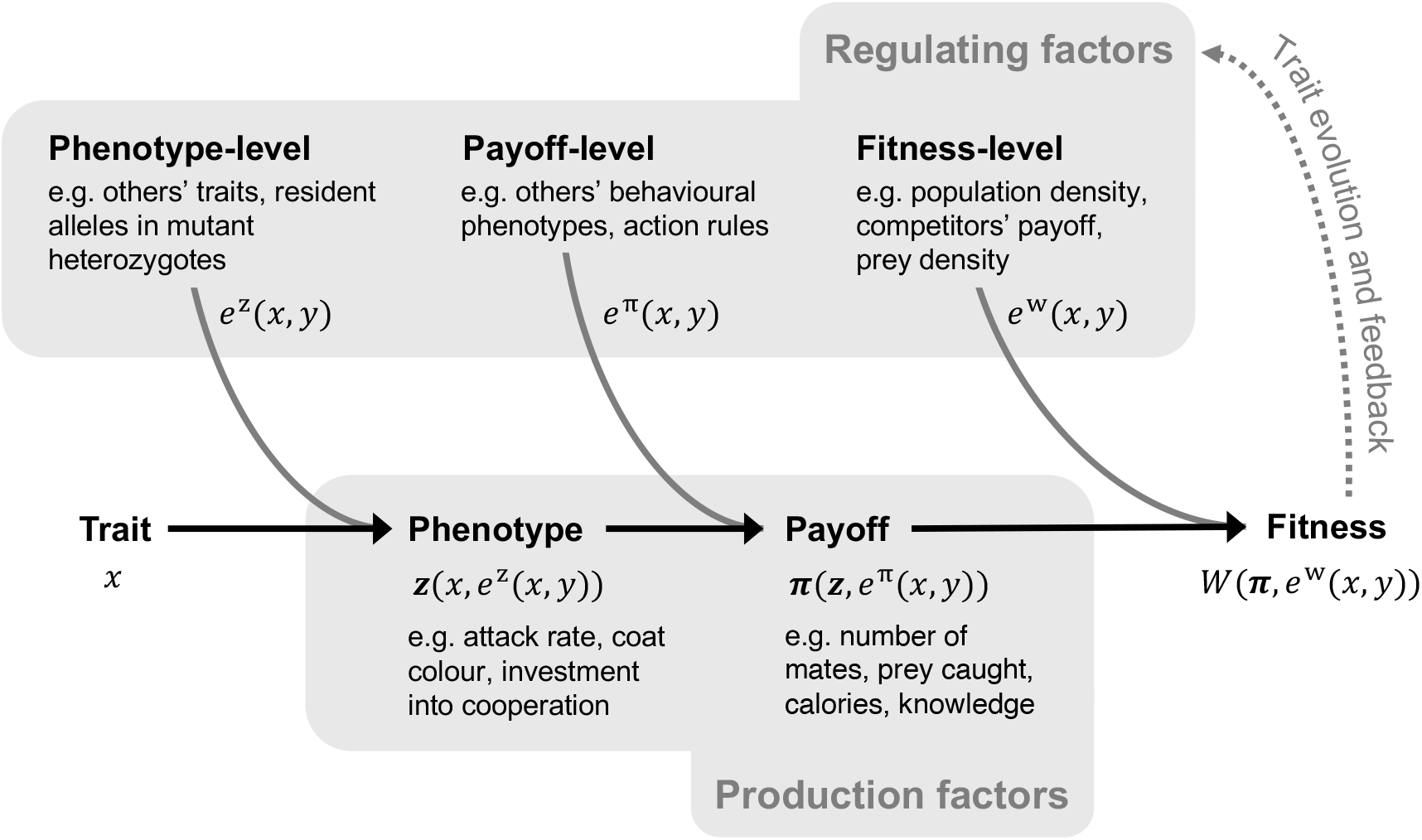
Production and regulating fitness factors. Diagram showing the construction of invasion fitness, from genetic trait value *x* through production factors: phenotype and payoff. These various layers and their regulating factors offer different opportunities for population feedback as trait evolution leads to changes in regulating factors. Section 3.2 for details.

### 3.3 Intensive versus extensive regulating factors and their trait dependence

To connect eq. (9) to the literature, it is useful to recognise that a regulating factor can be one of two types. The first type a regulating factor can be is an extensive variable, which is a variable indicating the size of a system under focus (Kondepuli and Prigogine, 2015), such as population size, number of competitors, or total resource availability. In evolutionary ecology, such variables are often treated as state variables whose equilibrium values are determined by the demographic structure or dynamics of the resident population (e.g. Diekmann et al., 2003; Pásztor et al., 2016). We illustrate this notion in Box 2, where regulating factors are determined by resident equilibrium equations (see also Diekmann et al., 2003, eq. 2.1; Kisdi and Geritz, 2016, eq. 8).

Alternatively, a regulating factor can be an intensive variable, which is a variable that is not indicative of the size of the system, i.e. variables that do not scale with system size (Kondepuli and Prigogine, 2015). Instead, these reflect per capita quantities, such as average survival, average fecundity, or local competitive pressure. This perspective is particularly common in population and quantitative genetics, or evolutionary game theory models with constant population size, where fitness is defined in terms of relative success (e.g., ratio of fecundities). This is the case for instance in the classical Wright-Fisher and Moran models of population genetics, where an individual’s fitness is expressed directly as a function of the survival probability and/or fecundity of others (e.g. Maynard Smith, 1982; Gillespie, 2004; Hartl and Clark, 2007; Ewens, 2004; Lynch and Walsh, 2018).

The expression of invasion fitness in eq. (9) in terms of regulating factors shows that mutant and resident traits can interact in many different ways, and hence that *I*(*y*) in eq. (6) is generally non-zero. Because fitness results from a hierarchy of nested interactions (trait → phenotype → payoff → fitness), *x* and *y* can influence *ρ*(*x, y*) through multiple biological pathways. The structure of eq. (9) allows us to identify three main pathways leading to NTDS: when interactions are (i) ecologically, (ii) socially, or (iii) genetically mediated, which we explore in turn below. For each, we characterise *I*(*y*) and illustrate how NTDS may arise through examples. Although these examples are based on scenarios that have been studied previously, all the polymorphisms we report here are new. In cases where selection leads to evolutionary branching, we also analyse the stability of the resulting oligomorphism for completeness in the Appendix (following the analysis detailed in Appendix C).

## 4 Ecologically mediated interactions

### 4.1 Ecological setup

Many interactions among individuals are indirect, occurring through environmental change, e.g., through resource consumption or habitat modification. In such cases, trait dependence arises via ecological feedbacks, where changes in population trait values alter environmental conditions, which in turn impact individual reproduction and survival. To formalise this, we set ***π***(**z**(*x*, **e**^z^(*x, y*)), **e**^*π*^(*x, y*)) = **z**(*x*) and **e**^w^(*x, y*) = **e**(*y*) in eq. (9) so that an individual’s payoff depends only on its own trait *x*, while the environment is determined solely by the resident trait *y*. Invasion fitness then simplifies to *ρ*(*x, y*) = *W* (**z**(*x*), **e**(*y*)). We refer to “ecologically mediated interactions” as cases where fitness takes this form, with interactions mediated through a regulating factor **e**(*y*) exogenous to the individual. These interactions can nevertheless involve complex ecological feedback loops, potentially spanning multiple environmental state variables (e.g., multitrophic consumer-resource models). In the ecological literature, the production factors **z**(*x*) are sometimes referred to as environmentally dependent parameters, while regulating factors **e**(*y*) are called competition parameters (Chesson, 2000a, p. 212-213).

For simplicity, we further assume that

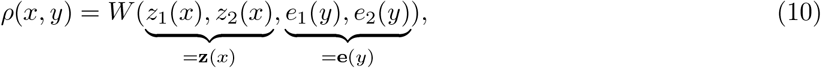

so that the function *W* depends on only four arguments, which are sufficient to capture standard pathways to trait-dependent selection under ecological interactions. The first two arguments, *z*_1_(*x*) and *z*_2_(*x*), are production factors, assumed to never decrease fitness such as fecundity or survival (*∂W/∂z*_1_ ≥ 0 and *∂W/∂z*_2_ ≥ 0). The second two, *e*_1_(*y*) and *e*_2_(*y*), are regulating factors, which are assumed to only feedback negatively on fitness (*∂W/∂e*_1_ ≤ 0 and *∂W/∂e*_2_ ≤ 0). As discussed in section 3.3, the regulating factors can be conceptualized as intensive variables, in which case one can set *e*_1_(*y*) = *z*_1_(*y*) and *e*_2_(*y*) = *z*_2_(*y*), or as extensive variables, which would then satisfy an equilibrium system of equations as detailed in Box 2.

Substituting eq. (10) into eq. (7a), shows that any singularity *y*\* must satisfy

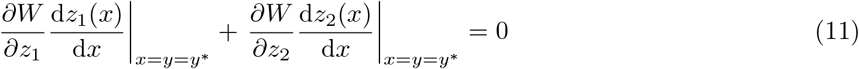

(here and throughout, we drop the argument of functions with multiple arguments when the notation becomes too cumbersome and there is no risk of confusion). Eq. (11)> shows that for a singularity *y*\* to exist, there must be a trade-off between the fitness returns obtained via the two production factors. Because fitness increases with both production factors (*∂W/∂z*_1_ ≥ 0 and *∂W/∂z*_2_ ≥ 0), selection would favour ever larger trait values if an increase in the trait raises both factors simultaneously. In that case, trait evolution would either be open ended or end up on a boundary of the trait space rather than at a singular point. An interior singularity therefore requires that changes in the trait increase one production factor while reducing the other, so that their fitness effects balance at *y*\*.

Because invasion fitness in a monomorphic population must equal one, i.e. *W* (*z*_1_(*y*), *z*_2_(*y*), *e*_1_(*y*), *e*_2_(*y*)) = 1, we also have

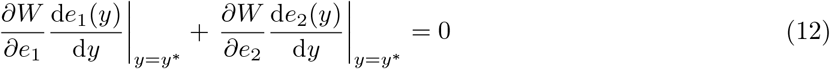

(using eq. 11). Since *∂W/∂e*_1_ ≤ 0 and *∂W/∂e*_2_ ≤ 0, eq. (12) indicates that d*e*_1_(*y*)*/* d*y* and d*e*_2_(*y*)*/* d*y* must have opposite signs at *y* = *y*\*. In other words, a trait increase in the population must be associated with opposite effects on regulating factors 1 and 2. This reflects a fundamental ecological or demographic constraint that arises when there is more than one regulating factor owing to density-dependent regulation: at a singularity, an increase in competition for one resource must be offset by a decrease in competition for another.

### 4.2 Interaction coefficient

We now turn to specifying the interaction coefficient *I*(*y*\*) under the above setup. Substituting eq. (10) into eq. (6) and using eqs. (11)–(12), we obtain

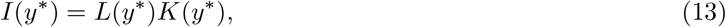

where *L*(*y*\*) = (d*z*_1_(*y*)*/* d*y*)(d*e*_1_(*y*)*/* d*y*)|_*y*=*y*_∗ is the product of the marginal changes in the first production and regulating factors; and

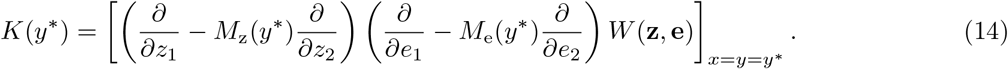

Here, *M*_z_(*y*\*) and *M*_e_(*y*\*) are marginal rates of factor substitution (Jehle and Reny, 2011, p. 128) (see Appendix D.1 for calculations). Specifically, *M*_z_(*y*\*) = (*∂W/∂z*_1_) */*(*∂W/∂z*_2_)|_*x*=*y*=*y*_∗ ≥ 0 is the rate at which production factor *z*_2_ can be substituted for *z*_1_ without changing fitness: it measures the change in the number of units of *z*_2_ required to compensate for a one-unit increase in *z*_1_ in terms of fitness effects. Similarly, *M*_e_(*y*\*) = (*∂W/∂e*_1_)*/*(*∂W/∂e*_2_)|_*x*=*y*=*y*_∗ ≥ 0 is the rate at which limiting factor *e*_2_ can be substituted for *e*_1_ without changing fitness.

Before interpreting *K*(*y*\*), let us remark that for most ecologically relevant models, *L*(*y*\*) will be nonzero (*L*(*y*\*) ≠ 0) since in the presence of a trade-off, neither a production factor nor a regulating factor will be pushed to a maximum by evolution (i.e. d*z*_1_(*y*)*/* d*y* ≠ 0 and d*e*_1_(*y*)*/* d*y* ≠ 0 at *y* = *y*\*). In fact, for any given model one should be able to define fitness factors such that *L*(*y*\*) *>* 0, since when a trait increases the production factor for one individual (e.g., a trait increases its fecundity), this is likely to increase a regulating factor when expressed by others (e.g., greater density of offspring before density-dependent competition for recruitment). In particular, when regulating factors are taken as intensive variables (such that *e*_1_(*y*) = *z*_1_(*y*), *e*_2_(*y*) = *z*_2_(*y*)), this leads to *L*(*y*\*) = (d*z*_1_(*y*\*)*/* d*y*\*)^2^ ≥ 0. Either way, condition (8) simplifies to

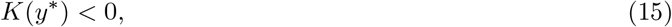

which therefore provides the condition for NTDS.

Eq. (14) can then be interpreted as follows. The first operator, (*∂/∂z*_1_ − *M*_z_(*y*\*) *∂/∂z*_2_), can be seen as measuring the net effect of a marginal change in the mutant trait *x* via the production factors, taking into account the fact that near *y*\*, an increase in one production factor necessarily entails a reduction in the other (recall eq. 11). This trade-off is captured by the coefficient −*M*_z_(*y*\*). The second operator, (*∂/∂e*_1_ − *M*_e_(*y*\*) *∂/∂e*_2_), measures the net effect of a marginal change in the population trait *y* via the regulating factors, taking into account the constraint that at resident demographic equilibrium, an increase in one regulating factor is necessarily associated with a reduction in the other (recall eq. 12). This constraint is captured by the coefficient −*M*_e_(*y*\*). When applied successively to *W* (**z, e**), the two operators thus quantify the effect of the interaction between changes in mutant and population trait on mutant fitness, as expected.

The structure of eq. (14) helps understand the origin of NTDS from ecologically mediated interactions. It shows that NTDS arises when the trait-induced trade-off in production factors (first operator) interacts negatively with the trait-induced constraint on ecological feedback (second operator). This can be illustrated with a local adaptation scenario (e.g. Levene, 1953; Gillespie, 1991; Kisdi and Geritz, 1999; Svardal et al., 2015). Suppose *z*_1_ and *z*_2_ are fecundity in two habitats (say a wet and a dry patch), while *e*_1_ and *e*_2_ are the corresponding local densities. Suppose also that a mutant increasing its trait value increases its fecundity under wet conditions, which by trade-off, necessarily means decreased fecundity in the dry patch. At *y*\*, these two direct effects are balanced. If at the same time, the population trait varies in the opposite direction, then density increases in the dry patch and decreases in the wet one. If this ecological feedback leaves the relative marginal value of fecundity in the two habitats unchanged, the production trade-off remains balanced and *K*(*y*\*) = 0. If by contrast, the decrease in wet-patch density and increase in dry-patch density make wet-patch fecundity more valuable relative to dry-patch fecundity, then the mutant change towards the wet patch becomes beneficial: the gain in the wet patch outweighs the loss in the dry one. This gives *K*(*y*\*) *<* 0, enabling NTDS and hence mutual invasibility (indeed, the same reasoning applies when the mutant changes to adapt to the dry habitat while the resident adapts to the wet one).

A direct consequence of eq. (13) is that polymorphism cannot emerge when invasion fitness depends on a single production or a single regulating factor, since in this case, *K*(*y*\*) = 0 and the necessary condition for polymorphism can never be met. Specifically, when only one production (regulating) factor, say *z*_2_ (*e*_2_) affects fitness while the other *z*_1_ (*e*_1_) does not, it follows that *M*_z_(*y*\*) = 0 (*M*_e_(*y*\*) = 0) and *∂*^2^*W/*(*∂z*_1_*∂e*_1_) = *∂*^2^*W/*(*∂z*_1_*∂e*_2_) = 0 [*∂*^2^*W/*(*∂z*_1_*∂e*_1_) = *∂*^2^*W/*(*∂z*_2_*∂e*_1_) = 0]. For regulating factors, this restates a classical result in theoretical ecology: at least two independent regulating factors are needed to sustain interspecific coexistence (e.g. Levin, 1970; Haigh and Maynard Smith, 1972; Meszéna et al., 2006; Pásztor et al., 2016). Eq. (14) additionally shows that,as established in Metz et al. (2008), an analogous requirement holds for production factors in the intraspecific case: polymorphism requires at least two production factors.

In addition to aiding interpretation, eq. (14) is also methodologically useful. Importantly, eq. (14) does not rely on an explicit specification of how traits influence fitness factors (i.e. how *x* and *y* influence **z** and **e**). This generality makes it valuable as it provides a direct means to identify whether polymorphism can be promoted in a given biological scenario. In the next sections, we apply this approach to two different scenarios, each revealing previously undescribed polymorphisms.

### 4.3 Life-history evolution and limited dispersal

We first consider a classic scenario of life-history evolution involving a trade-off between fecundity and survival, two fundamental production factors (e.g. Charnov, 1997; Case, 2000). Although such trade-offs have been studied extensively, it remains unresolved whether polymorphism can emerge when dispersal is limited and generations overlap. To address this, we consider a subdivided iteroparous population with the following life-cycle events: (a) Each adult individual occupies a site where it produces a large number of juveniles and then either survives or dies. (b) Each juvenile either migrates out of the natal site with probability *m* or with probability 1 − *m* remains philopatric.^3^ (c) Density-dependent competition occurs locally among juveniles for the breeding spots vacated by adult mortality, so that the adult population is regulated back to a constant level. With *z*_1_(*x*) denoting the survival probability of an adult expressing trait *x* and *z*_2_(*x*) its fecundity, invasion fitness for this model is

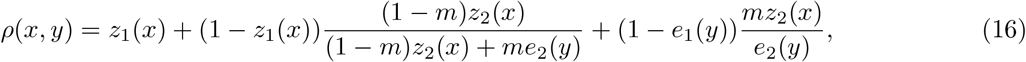

where the regulating factors are *e*_1_(*y*) = *z*_1_(*y*) and *e*_2_(*y*) = *z*_2_(*y*), i.e. the resident’s survival and fecundity (intensive variables). With complete dispersal *m* = 1, eq. (16) reduces to the standard reproductive-effort model in panmictic populations, for which polymorphism cannot emerge (Charnov, 1997; Case, 2000). More generally, eq. (16) coincides with the model of Pen (2000) for one individual per patch and to the model of Ronce and Promislow (2010) in the absence of senescence; however, those studies focused on directional selection and did not analyse the emergence of polymorphism.

Under model (16), the singular reproductive effort *y*\* involves a positive allocation to reproduction, and this allocation increases as dispersal becomes more limited (Pen, 2000, eq. 8; see also eq. A-12 in Appendix here). This reflects a trade-off between survival and reproduction: survival secures the natal breeding site for a mutant lineage, whereas reproduction increases the number of offspring competing for other sites. When dispersal is limited, fewer competitors arrive at the natal site, so it is likely to remain within the lineage even if the adult does not survive. As a result, the selective advantage of survival declines relative to reproduction, favouring greater reproductive effort. What remains to be determined is whether selection is negatively trait-dependent at this singularity, thereby opening the door to the emergence of polymorphism.

Substituting eq. (16) into eq. (14) and using eq. (11), we obtain

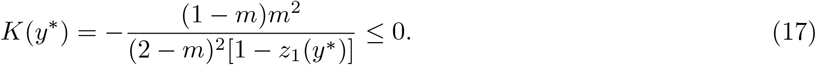

This shows that NTDS arises only under partial dispersal 0 *< m <* 1. To understand why, it is useful to consider that the mutant allocates more to reproduction (and less to survival), while individuals in the population allocate more to survival (and less to reproduction). The mutant’s change increases the number of juveniles it produces, whereas the change in the population trait reduces the number of breeding sites available. When dispersal is complete, these two effects cancel one another, so that the gain in mutant fecundity is balanced by the reduction in available breeding sites caused by the population change. When dispersal is partial, however, this balance breaks down because lower resident fecundity also reduces the number of immigrant juveniles arriving at the mutant’s natal site. If the mutant adult dies, its philopatric offspring therefore face weaker competition for the local vacancy. As a result, mutant invasion fitness increases when its trait diverges from that of the resident. This NTDS should enable the coexistence of different life-history strategies, given an appropriate trade-off between survival and reproduction.

We investigated this specifically with a trade off induced by *z*_1_(*x*) = *s*_0_*x*^*α*^ and *z*_2_(*x*) = (1 − *x*)^*β*^ with parameters *α <* 1, *β <* 1 and *s*_0_ *<* 1. Our analysis in Appendix D.2 shows that, for a range of parameter values, polymorphism in resource allocation can then be favoured by gradual evolution as long as 0 *< m <* 1. The end-point of evolution consists of two morphs using mixed allocation to survival and fecundity, with the most abundant morph investing more in survival. We conducted individual-based simulations that confirm these findings (Fig. 2). As *s*_0_ ≈ 1, i.e. which can be thought of as a situation with almost no external mortality, one morph invests essentially all resources in survival becoming quasi immortal (see Appendix D.2 for more details). The population can thus display a marked dimorphism in life-history strategies (see Appendix D.2 for more details), a dimorphism impossible under well-mixed conditions (Charnov, 1997; Case, 2000).

**Figure 2.**
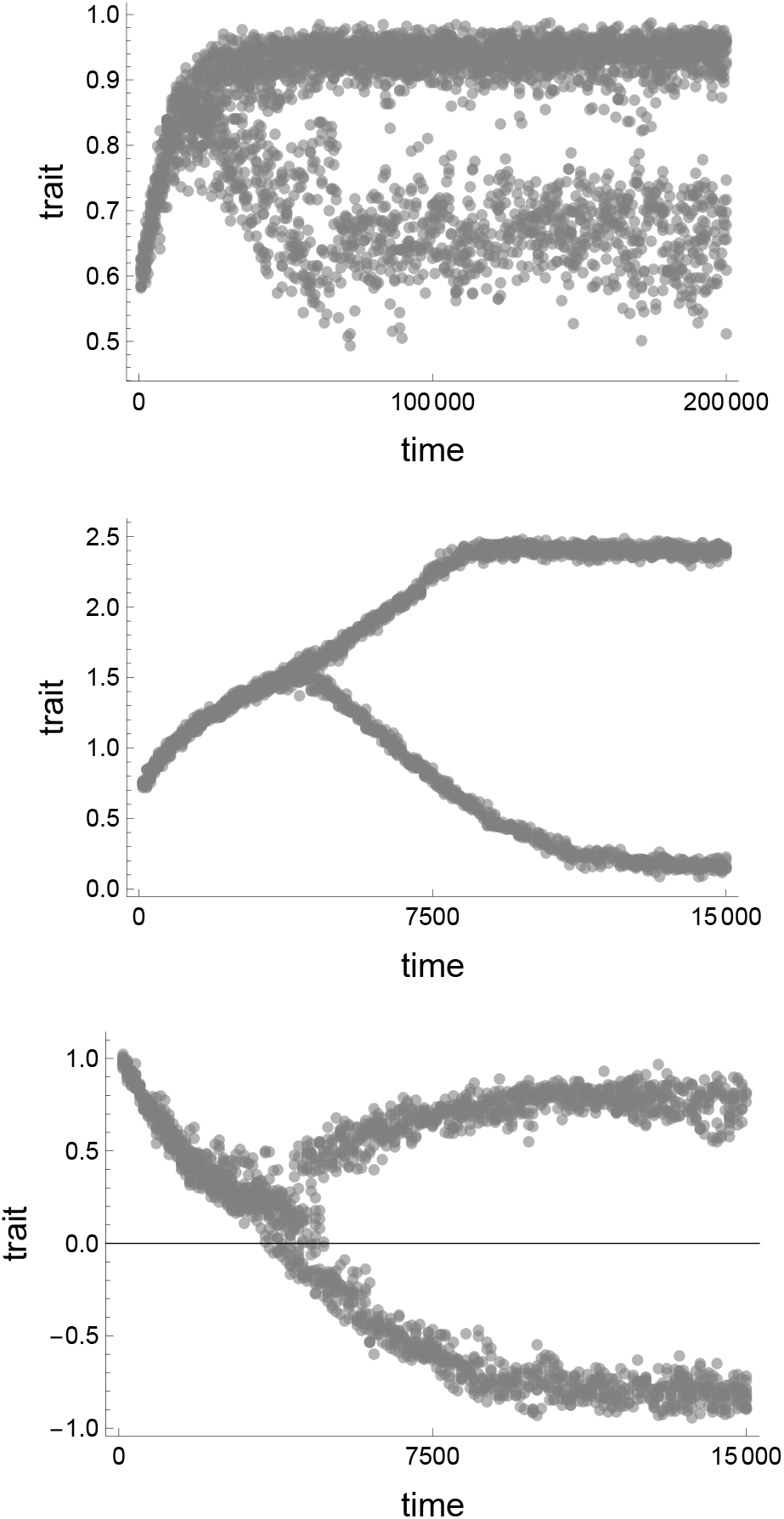
Illustration of evolutionary branching and long term evolution under individual-based simulations for a sample of novel polymorphisms. Top panel is for the fitness model eq. (16), which leads to the coexistence of a quasi-immortal morph investing most resources into survival and another one expressing a more balanced mixed strategy (the trait *y* is here investment into survival, with fraction 1 − *y* invested into fecundity, see Appendix D.2 for the explicit fitness factors used, the formal analysis, and parameter values). Middle panel is for payoff model eq. (28), which leads to the coexistence of one morph investing a lot into contests (“Hawk”) and another one investing moderately (“Dove”) (the trait *y* here is investment into contest, see Appendix E.2.1 for the fitness factors used, the formal analysis, and parameter values). Bottom panel is for the payoff model eqs. (31)–(33), which leads to the coexisting of optimists and pessimists (the trait *y* here is a self esteem bias, with *y >* 0 describing optimism and *y <* 0 pessimism, see Appendix E.2.2 for the formal analysis and parameter values). In each case, reproduction follows a Wright-Fisher process (lottery model) with *N* = 10000 individuals. Mutations occur with probability 0.015 and effect sizes are sampled from a Normal distribution with mean 0 and standard deviation 0.01 (trait values are capped between zero and one in the top panel).

### 4.4 Temporal heterogeneity and limited dispersal

The second scenario we examine is temporal heterogeneity, where production factors vary between generations, which is a classical source of diversity in both evolutionary ecology and population genetics (e.g. Chesson and Warner, 1981; Gillespie, 1991; Turelli et al., 2001). Gradual evolution under temporal heterogeneity and limited dispersal has previously been analysed in models assuming infinitely large local populations, effectively excluding kin competition (Svardal et al., 2015). Here we consider the complementary case of populations with limited dispersal and one individual per site, where kin competition arises under hard selection.

Consider again a population subject to limited dispersal, and that experiences two environmental states, say a wet and a dry environment, occurring independently in each generation with probabilities *q* and 1 − *q*, respectively. At each demographic time step, each adult individual occupies a site where it produces a large number of juveniles, with fecundity *z*_*i*_(*x*) depending on both its trait *x* and the environmental state *i* ∈ {1, 2}. Adults then either survive with probability *s* or die, freeing up breeding spots. Each juvenile either disperses with probability *m* or remains philopatric. Density-dependent competition among juveniles occurs locally for the breeding spots vacated by adult mortality. Invasion fitness for this scenario then takes the form of a geometric mean across environmental states, i.e.

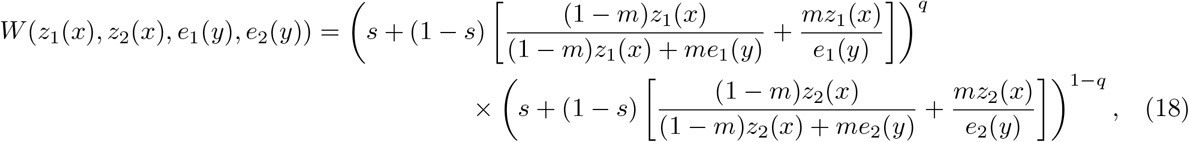

where *e*_1_(*y*) = *z*_1_(*y*) and *e*_2_(*y*) = *z*_2_(*y*).

Substituting eq. (18) into eq. (14) and eq. (11) yields

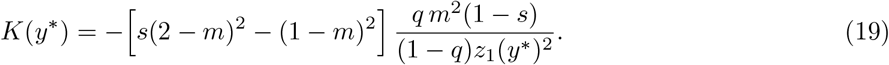

Whether *K*(*y*\*) *<* 0 depends on survival *s* and dispersal *m*. Consider first the case of complete dispersal (*m* = 1). When generations do not overlap (*s* = 0), *K*(*y*\*) = 0 and polymorphism is impossible, whereas when generations overlap (*s >* 0), *K*(*y*\*) *<* 0 and polymorphism becomes possible. To understand this, consider a mutant and a population whose traits change in opposite directions from *y*\*, so that the mutant performs better in one environment and resident individuals in the other. When *s* = 0, the fitness loss experienced by the mutant in one environment is offset by the gain in the other. When generations overlap (*s >* 0), this balance changes due to a storage effect (Chesson, 2000b): survival carries individuals across years, so the composition of the surviving adult pool reflects which trait values were favoured previously. When environments differ in which trait values they favour, increasing the performance of the resident population in a given environment builds up the representation of that trait among survivors, thereby increasing effective competition in that environment. Consequently, when mutant and population traits change in opposite directions, the mutant tends to experience weaker competition in the environment where it performs best. This yields NTDS.

Eq. (19) shows that this effect weakens as dispersal decreases, with *s >* (1 − *m*)^2^*/*(2 − *m*)^2^ required for NTDS to occur. Survival and limited dispersal thus have opposing effects on NTDS. While overlapping generations promote NTDS via a storage effect (Chesson, 2000b), limited dispersal, by contrast, increases local kin competition, so that offspring are more likely to compete against individuals sharing the same trait. This reduces the advantage of a mutant performing well in a different environment from the resident population and erodes the asymmetry that drives negative trait-dependent selection.

Using *z*_*i*_(*x*) = exp(−(*x* − *o*_*i*_)^2^*/*(2*σ*^2^)), it can readily be shown that with fitness eq. (18) there exists a range of parameter values leading to evolutionary branching, and that this range becomes smaller as dispersal becomes limited (see Appendix D.3). This differs from the results of Svardal et al. (2015), where dispersal affected diversification only under spatial heterogeneity (i.e. their eq. 15), and had no influence under purely temporal heterogeneity with effectively infinite local population size (and hence no kin competition). In our model, by contrast, limited dispersal generates kin competition, which weakens NTDS and inhibits branching, consistent with previous results showing that kin competition tends to oppose diversification under constant environments (Day, 2001; Ajar, 2003). The role of kin selection in polymorphism is explored further in section 6.2.

## 5 Socially mediated interactions

We now turn to another important class of trait-dependent interactions: those mediated by direct social interactions influencing payoffs. Classic examples include contests for mates or territories, public-good investments, cooperative breeding, and various forms of aggressive or cooperative encounters (e.g. Maynard Smith, 1982; Vincent and Brown, 2005; McNamara and Leimar, 2020). Here, an individual’s payoff explicitly depends on the traits expressed by others in the population, creating a social feedback loop that may lead to stable polymorphisms.

### 5.1 Selection and interaction coefficients at the payoff level

To capture socially mediated interactions in a simple way covering multiple scenarios, we assume that the payoff to an individual is one-dimensional, given by ***π***(**z**(*x*, **e**^z^(*x, y*)), **e**^*π*^(*x, y*)) = *π*(*x, y*), and depends explicitly on both mutant and resident traits. Additionally, we assume that the regulating variable **e**^w^(*x, y*) = *e*(*y*) is also one dimensional and depends solely on the resident population. Under these assumptions, invasion fitness (9) takes the form

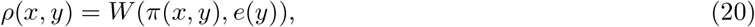

for some mapping *W*. We assume that invasion fitness is monotonically increasing with payoff (i.e. *W* increases with its first argument). Eq. (20) represents the standard fitness formulation that is implicitly used in evolutionary game theory, encompassing both pairwise interactions and cases where individuals interact collectively (“playing against the field”; e.g. Maynard Smith, 1982; Vincent and Brown, 2005; McNamara and Leimar, 2020). This formulation captures the essence of socially mediated interactions, where fitness depends explicitly on payoffs (determined by the traits of interacting individuals), while environmental factors influence selection only indirectly through their effect on population-wide competition or regulation.

Substituting eq. (20) into eq. (6), we show in Appendix E.1 that the invasion analysis can be fully worked out from the payoff function. In particular, any singular strategy *y*\* must be such that

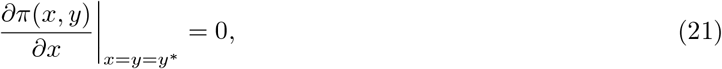

and the necessary condition for protected polymorphism (eq. 8) reduces to

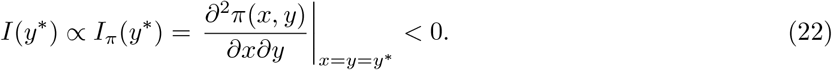

When condition (22) holds, this indicates that the mutant’s payoff increases when the mutant and the resident population change trait values in opposite ways. Since fitness is monotonically increasing with payoff, these payoff effects directly generate NTDS. We explore further how such selection can emerge owing to social interactions by making more specific assumptions concerning payoff in the next section.

### 5.2 Action based feedback

To investigate how a social feedback can promote polymorphism, we consider a class of models in which an individual’s payoff depends explicitly on actions taken during interactions. Specifically, we assume that the payoff can be expressed as:

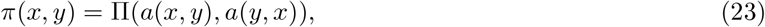

for a mapping Π, where *a*(*x, y*) and *a*(*y, x*), are the actions of a mutant and a resident individual, respectively (formally actions here are equivalent to phenotypes *z* but for ease of interpretation in a social context, we will refer to actions and use the variable *a* in this section). These actions may depend not only on an individual’s own trait but also on the trait of its interaction partner.

This formulation covers two broad classes of interactions. The first is pairwise interactions, where *a*(*x, y*) and *a*(*y, x*) describe the actions of two individuals in a direct encounter. The dependence of an individual’s action on its own trait and that of its partner allows us to model cases where actions arise as equilibrium outcomes of a dynamic behavioural process, such as learning, information exchange, or negotiation (as used in models of evolving response rules, motivations, and preference evolution, e.g. McNamara et al., 1999; André and Day, 2007; Heifetz et al., 2007a; Akçay and Van Cleve, 2012; Alger and Lehmann, 2023). This also captures simpler pairwise interactions where one’s action depends only on own trait, *a*(*x, y*) = *a*(*x*), as is very commonly studied in evolutionary game theory (Maynard Smith, 1982; McNamara and Leimar, 2020). In any case, the trait in the resident population *y* appearing in *a*(*x, y*) corresponds to the regulating factor **e**^z^(*x, y*) = *y* acting at the level of the phenotype (i.e. the action), while the action *a*(*y, x*) corresponds to the regulating factor **e**^*π*^(*x, y*) = *a*(*y, x*) at the level of payoff (recall eq. 9).

The second class of interactions covered by eq. (23) corresponds to “playing against the field” multiplayer interactions (Maynard Smith, 1982). In this case, actions depend only on an individual’s own trait, such that *a*(*x, y*) = *a*(*x*) and *a*(*y, x*) = *a*(*y*). This extends beyond pairwise interactions, as multiple individuals expressing action *a*(*y*) can simultaneously influence the individual’s payoff. Here, there is no regulating factor at phenotype level, and the regulating factor at the payoff level simplifies to **e**^*π*^(*x, y*) = *a*(*y*).

Substituting eq. (23) into eq. (21) reveals that any singularity *y*\* must be such that

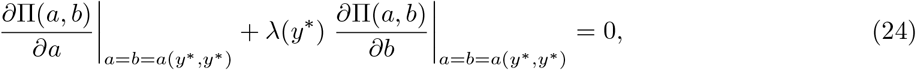

where

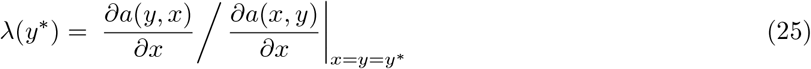

is a response coefficient, which measures how much the partner to a mutant changes her action in response to a change in the action of the mutant (a change that is due to a trait change in the mutant). When *λ*(*y*\*) *>* 0, an increase in one’s action due to a trait change leads to a corresponding increase in the partner’s action. Conversely, when *λ*(*y*\*) *<* 0, the response is negative, meaning that a trait-driven increase in one’s action is associated with a decrease in the partner’s action. Eq. (24) expresses that at equilibrium *y*\*, selection balances two opposing effects on payoff: a direct effect due to changes in the focal mutant’s own action (first term) and an indirect effect due to the response of the partner’s action (second term), weighted by the response coefficient *λ*(*y*\*).

The balance captured by eq. (24) helps to understand the evolution of many well-known social interactions. For instance, when payoff Π(*a, b*) is decreasing in its first argument and increasing in an accelerating manner in its second argument, individual actions can be regarded as levels of cooperation. If actions increase with trait values, and if *a*(*x, y*) = *a*(*x*) and *a*(*y, x*) = *a*(*y*) so that individuals cannot respond to their partner’s action, then *λ*(*y*\*) = 0. In this case, no singularity exists, and eq. (24) correctly predicts the loss of unconditional cooperation in a well-mixed population. By contrast, when cooperation is conditional, so that actions depend on both players’ traits enabling individuals who express cooperative behaviour to receive cooperative responses in return, then conditional cooperation via reciprocity can evolve when *λ*(*y*\*) *>* 0. This occurs for instance where traits determine response slopes such that equilibrium actions are given by *a*(*x, y*) = *α* + *xa*(*y, x*) and *a*(*y, x*) = *α* + *ya*(*x, y*) for some *α >* 0 (e.g. McNamara et al., 1999; Taylor and Day, 2004). More broadly, the coefficient *λ*(*y*\*) plays a central role across many scenarios of social evolution (e.g. Akçay and Van Cleve, 2012; Akçay and Cleve, 2014; Lehmann, 2022; Alger and Lehmann, 2023).

Using eq. (24) and eq. (25) on substituting eq. (23) into eq. (22), we find that *I*_*π*_(*y*) = *L*_*π*_(*y*\*)[*K*_*π*_(*y*\*)+*K*_*λ*_(*y*\*)], where *L* _*π*_ (*y*\*) = (*∂a*(*x, y*)*/∂x*)^2^|_*x*=*y*=*y*_* ≥ 0, such that NTDS selection occurs when

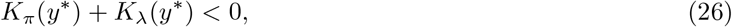

in which

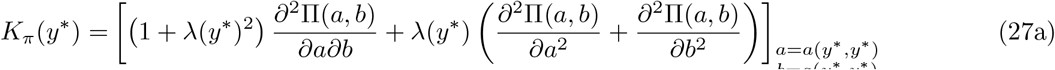

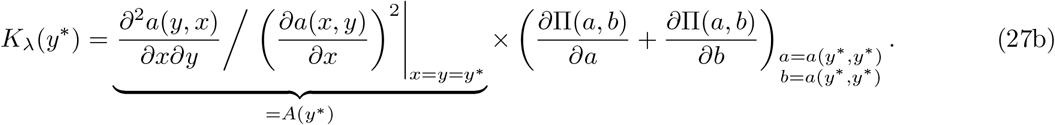

The first component, *K*_*π*_(*y*\*), describes trait-dependent selection via direct payoff effects, which considers how selection on the mutant changes as the population trait is varied, while keeping the response coefficient *λ*(*y*\*) unchanged. This induces NTDS through two pathways. First, NTDS occurs where the effects of actions on payoff are antagonistic, meaning that the actions of interacting individuals interfere in a way that reduces payoff, i.e. *∂*^2^Π(*a, b*)*/*(*∂a∂b*) *<* 0. When this holds, a mutant obtains greater payoffs when the actions of interacting individuals change oppositely to the mutant. Second, NTDS can occur where the payoff functions change non-linearly with the actions of interacting individuals. In particular, NTDS will be favoured when payoff accelerates with both actions (i.e. *∂*^2^Π*/∂a*^2^ *>* 0 and *∂*^2^Π*/∂b*^2^ *>* 0) and if the response coefficient is negative, *λ*(*y*\*) *<* 0, meaning that the action of the mutant and that of its interaction partner vary in opposite directions at a singular point. When this is the case, a mutant individual who increases its action will, by eliciting a reduction in the action of its partner, have a higher payoff owing to these non-linearities.

The second component in eq. (27), *K*_*λ*_(*y*\*), describes trait-dependent selection via response effects, i.e. through changes in *λ*(*y*). This is governed by *A*(*y*\*), which measures how one’s action changes in response to a similar trait change in others. When *A*(*y*\*) *<* 0 changing traits similarly reduces the response coefficient, whereas when *A*(*y*\*) *>* 0 it increases it. For example, if *∂*Π*/∂a* + *∂*Π*/∂b >* 0, as occurs by definition for cooperative actions (*sensu* the social semantics of Rousset, 2004; Bshary, 2007) and *A*(*y*\*) *<* 0 (for instance, due to interference in cooperative investments) an increase in population trait leads to weaker cooperation overall, thus favouring NTDS. We next illustrate how eq. (26) helps understand two more specific examples of social interactions generating NTDS.

### 5.3 Examples

#### 5.3.1 Contests over resources

Consider a scenario where payoffs are determined by contest competition among *N* individuals for access to a resource of value *v*. Suppose the payoff function is

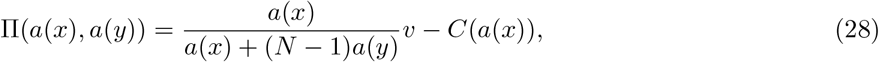

where the action *a*(*x*) measures the fighting or competitive effort of an individual with trait *x* and *C*(*a*(*x*)) is the cost associated with such effort. The probability of winning the contest depends on an individual’s relative effort, given by the ratio *a*(*x*)*/*[*a*(*x*) +(*N* − 1)*a*(*y*)], which is standard in conflict and contest competition (e.g. Hirshleifer, 1987; Skaperdas, 1996; Cant, 2012). This is a scenario of “playing against the field”, where actions depend only on one’s own trait and not on the trait of interacting partners so that the response coefficient (eq. 25) is zero *λ*(*y*) = 0, which implies *A*(*y*\*) = 0 and as a result, *K*_*λ*_(*y*\*) = 0.

Whether trait-dependent selection is negative therefore depends solely on whether actions are strategic substitutes at the payoff level (from eq. 27), which using eq. (28) yields

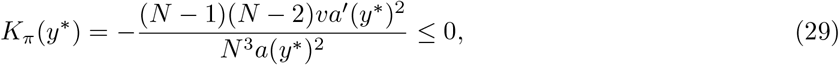

where the prime denotes the derivative. This equation shows that as long as *N >* 2 (meaning that at least three individuals compete for a resource), NTDS occurs. When *N* = 2, payoffs depend only on the direct outcome of the contest between two individuals, so the marginal effects of mutant and population trait changes combine additively and NTDS cannot occur. This contrasts with the classic Hawk-Dove game, where pairwise interactions (i.e. *N* = 2) can generate NTDS. In the Hawk-Dove game, payoffs decrease for aggressive individuals when interacting with other aggressive individuals, meaning that NTDS arises from interaction effects on costs (this would correspond here to writing the cost function as *C*(*x, y*) with *∂*^2^*C*(*x, y*)*/*(*∂x∂y*) *>* 0, where the cost of aggression increases as the opponent’s aggression increases). Without such interaction effects on costs, competition must involve multiple individuals for NTDS to emerge (*N >* 2). Nevertheless, the ease with which NTDS arises in our model exemplifies how conflicts in general provide a natural setting for polymorphism, since if one individual increases its competitive effort, this is likely to be mitigated if others in the population change similarly. In fact, evolutionary diversification has been found to occur through gradual evolution in multiple conflict models (McNamara and Leimar, 2020).

To see polymorphism emerge under fitness model (28), one can use *a*(*x*) = *e*^*ωx*^ and *C*(*a*(*x*)) = *x*^2^ with *ω >* 0 as a parameter. For *N >* 2, there exists a range of parameter values leading to evolutionary branching with these functions. Over the long term, gradual evolution results in a convergence stable and uninvadable dimorphism, with one morph investing heavily in competition while the other adopts a more moderate investment strategy. Further details on the analysis are provided in Appendix E.2.1, and Fig. 2 shows the emergence of this previously unreported polymorphism in individual-based simulations. This model with fixed number of contestants can readily be extended to a more realistic setting with random number of contestants *N* (independent of traits). In this case, NTDS requires that the expectation over the (random) number of contests yield E[(*N* − 1)(*N* − 2)*/N* ^3^] *>* 0.

#### 5.3.2 Evolution of preferences

The previous example had no behavioural response (*λ*(*y*\*) = 0); we now consider a case where individuals adjust their behaviour in response to their partner’s traits (*λ*(*y*\*) ≠ 0). We assume individuals interact in a pairwise manner and aim to maximise a utility function that evolves under a situation of complete information (see Alger, 2023 for discussion of information settings). This follows models of preference or motivation evolution, where selection acts not directly on the actions individuals take but on the underlying behavioral mechanism guiding their decision-making (e.g. Heifetz et al., 2007b; Akçay and Van Cleve, 2012; Alger and Lehmann, 2023; see Alger, 2023 for a review).

Building on Heifetz et al. (2007a,b), we assume that each individual’s utility consists of their own material payoff and a self-esteem bias, which distorts their perception of their own actions. Specifically, the utility functions for a mutant with action *a* and trait *x*, and a resident with action *b* and trait *y*, are given by

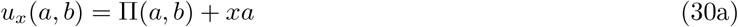

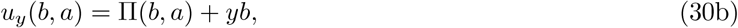

respectively. Here, the self-esteem bias is modulated by the evolving trait (*x* in mutants, *y* in residents) and determines whether individuals overestimate or underestimate the benefits of their own actions. When *x* = 0, a mutant individual simply seeks to maximise its material payoff. But when *x >* 0, the mutant overestimates the benefits of its own action, acting as if its actions are more rewarding than they actually are. Conversely, when *x <* 0, the mutant underestimates its returns and behaves more cautiously, even when a greater action value would be materially beneficial. In this way, the evolving trait *x* governs an individual’s “disposition” (Heifetz et al., 2007a,b), shaping whether they are overconfident optimists (*x >* 0) or underconfident pessimists (*x <* 0) in how they perceive and respond to interactions.

To complete the model, we assume that material payoffs follow a linear-quadratic structure, given by

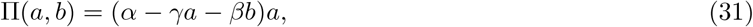

for parameters assumed to satisfy *α >* 0, *γ >* 0, and *β* ≥ *γ*. Eq. (31) is consistent with a model where actions constitute the effort or energy allocated to the extraction of a biotic resource that pairs of individuals exploit (e.g. pairs of individuals exploit different resource patches), and where the resource has logistic growth and a linear functional response to extraction. In this case, the parameters of eq. (31) can be interpreted as follows. The parameter *α* is proportional to the intrinsic growth rate of the resource and *γa* to the rate of resource extraction by an individual with action *a*. Assuming fast resource dynamics, the equilibrium amount of resource is then proportional to (*α* − *γa* − *γb*), and on re-scaling parameters^4^ the amount of resources individual with action *a* extracts per unit time is (*α* − *γa* − *γb*)*a*. Finally, we add to this *ab*(*γ* − *β*), which can be seen as a cost stemming from interference competition: when *β > γ*, joint efforts lead to diminishing returns, similarly to interference competition where competition over shared resources reduces overall resource uptake rate or hurts interacting individuals (e.g., Begon and Townsend, 2021, chapter 5).

By maximising their utility given the action of their partner, each individual plays a Nash equilibrium strategy. The necessary first-order conditions for a Nash equilibrium require that neither individual can improve their utility by unilaterally changing their action, meaning *∂u*_*x*_(*a, b*)*/∂a* = 0 and *∂u*_*y*_(*b, a*)*/∂b* = 0. Using eqs. (30) and (31), these conditions can be expressed as the system:

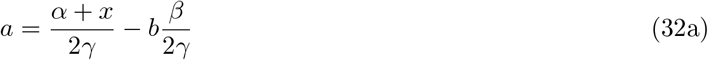

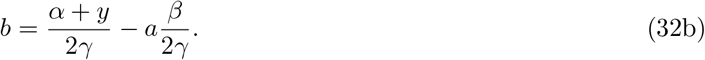

Solving these for *a* = *a*(*x, y*) and *b* = *a*(*y, x*) yields

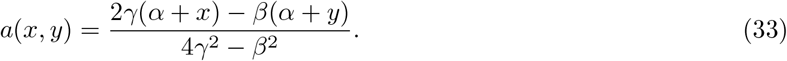

This solution represents a Nash equilibrium action because it is unique and satisfies the second-order condition for stability: *∂*^2^*u*_*x*_(*a, b*)*/∂a*^2^ = *∂*^2^Π(*a, b*)*/∂a*^2^ = −2*γ* ≤ 0 and *∂*^2^*u*_*y*_(*b, a*)*/∂b*^2^ = *∂*^2^Π(*b, a*)*/∂b*^2^ = −2*γ* ≤ 0. Eq. (33) shows that an individual’s action can increase with their own optimism (*x*) but decreases with that of their partner (*y*), provided 4*γ*^2^ *> β*^2^. The joint increase in actions by both players then reduces individual resource extraction rate when 2*γ > β > γ >* 0. To avoid these deleterious effects, it is best to reduce one’s own action when faced with more optimistic individuals (who increase their action as a result). This relationship is captured by the negative response coefficient *λ*(*y*\*) = −*β/*(2*γ*) (found by plugging eq. 33 in eq. 25), indicating that an increase in the partner’s action leads to a decrease in one’s own.

A key feature of eq. (32) is that an individual’s action is a linear response to their partner’s. This is formally identical to models for the evolution of response rules, where individuals adjust their behaviour based on social interactions (e.g., McNamara et al., 1999; Taylor and Day, 2004; see Alger, 2023 for a discussion on the connection between preference evolution and response rule evolution models). This linearity means that, here and in these aforementioned models, *A*(*y*) = 0 (eq. 27b). As a result, there can be no trait-dependent selection via a feedback on behaviour rules (i.e. *K*_*λ*_(*y*\*) = 0).

Therefore, NTDS arises when *K*_*π*_(*y*\*) *<* 0 in this model. Substituting eqs. (31) and (33) into eq. (27a), we obtain

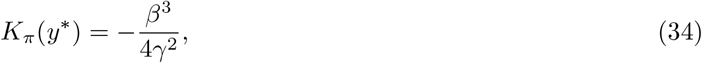

which is negative under our assumption of interference competition (*β > γ >* 0). It is then straightforward to show that evolutionary branching in this self-esteem trait can happen as long as 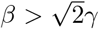. When it occurs, evolutionary branching leads to ever more “optimists” (with increasingly positive trait values) and ever more “pessimists” (with increasingly negative trait values) as both strategies fuel one another. This open ended evolution of exaggerated self-perception comes to a halt where there is a cost to overconfidence (Appendix E.2.2 for analysis and Fig. 2 for individual-based simulations).

The original study by Heifetz et al. (2007a,b) did not analyse the scope for polymorphism in self-esteem biases, and in fact considered parameter values generating evolutionarily stable singularities only. Heifetz et al. (2007a,b) also considered an alternative model of preference, in which the evolving trait *x* measures an individual’s other-regard. In this formulation, mutant and resident utilities take the form *u*_*x*_(*a, b*) = Π(*a, b*) + *x*Π(*b, a*) and *u*_*y*_(*b, a*) = Π(*b, a*) + *y*Π(*a, b*). We analyse this model in Appendix E.2.3 using eq. (31) for payoff. We show that for this model NTDS arises through payoff effects (*K*_*π*_(*y*\*) *<* 0) as well as via action response effects (*K*_*λ*_(*y*\*) *<* 0). Thus, polymorphism in social preferences can emerge in this model too (and see Cheikbossian and Peña, 2026 for similar findings in a model with another payoff function).

## 6 Genetically mediated interactions

Up until now, we have assumed that individuals are haploid. We now relax this assumption and investigate its implications for adaptive polymorphism. We will see that these depend on the nature of interactions between genes within individuals. We will also consider implications of the interaction between mutant copies in different individuals (hence “genetically mediated interactions”).

### 6.1 Diploidy and heterozygote advantage

We consider now that individuals are diploid and that an individual’s phenotype depends on the two homologous alleles it carries. To model this, we denote as *z*_*i*_(*x, y*) the phenotype *i* of an individual carrying a mutant allele with heritable trait value *x* and a resident allele with heritable trait value *y*. As such, an individual homozygote for the resident allelic value *y* expresses phenotype *z*_*i*_(*y, y*). We assume that mating is random so that the mutant allelic form *x* is only ever present in heterozygous state. We also assume that the genotype-to-phenotype map is: (i) homogeneous (by which we mean *z*_*i*_(*x, y*) = *z*_*i*_(*y, x*)) and (ii) such that variation in allelic value always leads to variation in phenotype, i.e. that phenotypes always show some degree of heritability (which entails *∂z*_*i*_(*x, y*)*/∂x*|_*x*=*y*_ = *∂z*_*i*_(*x, y*)*/∂y*|_*x*=*y*_ ≠ 0, where the first equality unfolds from assumption (i)). Where the phenotype of heterozygotes is a convex combination of the phenotypes of homozygotes, assumptions (i) and (ii) entail additive gene action (i.e. entail that *z*_*i*_(*x, y*) = *z*_*i*_(*x, x*)*/*2 + *z*_*i*_(*y, y*)*/*2); our forthcoming results apply more broadly (though the assumption that *∂z*_*i*_(*x, y*)*/∂x*|_*x*=*y*_ = *∂z*_*i*_(*x, y*)*/∂y*|_*x*=*y*_ amounts to local additivity).

#### 6.1.1 No effect of diploidy on the conditions for the emergence of polymorphism of a single phenotype

Let us first consider the case where invasion fitness depends on a single phenotype such that we can write

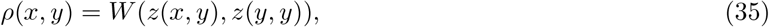

where the function *W* gives the invasion fitness of a *x/y* heterozygote in a population of *y* homozygotes. Substituting eq. (35) into eqs. (6)-(7a) and using our assumptions concerning *z*(*x, y*), we show in Appendix F.1 that the population converges to a singular allelic value *y*\*, which generates phenotype *z*\* = *z*(*y*\*, *y*\*), and subsequently undergoes evolutionary branching when the following holds:

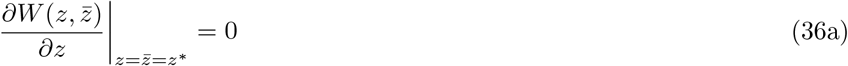

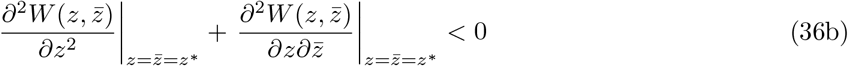

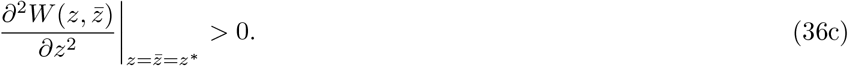

In these expressions, the derivatives are taken with respect to phenotype (where 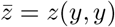 denotes resident phenotype) and not genetic values. These conditions are equivalent to those derived under haploidy (mirroring eq. 7a and conditions discussed thereafter). This shows that diploidy does not affect the conditions for the emergence of polymorphism via evolutionary branching, at least when underlying a single phenotype. In particular, the emergence of polymorphism requires that the cross derivative of the fitness function with respect to mutant and resident phenotypes is negative:

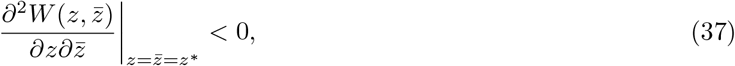

indicating that directional selection depends negatively on the phenotype expressed in the resident population (as in eq. 8 at the level of the trait value). Consequently, all our results in sections (4)–(5), which are based on an invasion fitness function of the form *ρ*(*x, y*), still apply unchanged to diploids, with *x* now standing for the phenotypic value of a mutant individual (*z*) and *y* for the phenotypic value in the resident population 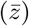.

Although the conditions favoring the emergence of polymorphism are left unchanged by diploidy under the above assumptions, diploidy does affect the condition for mutual invasibility (in fact, unlike the haploid case, it is possible to have mutual invasibility without NTDS owing to genetic constraints under diploidy, Appendix eq. A-43 for details). Additionally, evolution after the onset of evolutionary branching will likely differ between haploid and diploid populations owing to heterozygotes with intermediate phenotypes in the latter. In a haploid population, evolutionary divergence initially leads to the coexistence of two morphs. In sexual diploids, by contrast, mating between homozygotes for these diverged alleles creates a third, intermediate heterozygote. In scenarios with two niches (e.g. two patch types), evolution then leads one of the three resulting types–either one of the two homozygotes or the heterozygote–to be maladapted (which type evolves to be maladapted depends on model parameters; Kisdi and Geritz, 1999). As a result of the genetic constraint of additive effects, a proportion of the phenotypic polymorphism in the population is therefore non-adaptive. This in turn creates selection on the genetic architecture of the trait to reduce such genetic load, e.g. through the evolution of dominance (Van Dooren, 1999). More generally, genetic constraints that lead to a genetic load favour the evolution modifiers of recombination, epistasis, or gene duplication, that should eventually lead to a phenotypic polymorphism mirroring that seen in haploid asexuals, where such genetic constraints are absent.

#### 6.1.2 Heterozygote advantage when there are multiple phenotypes

Next, we consider invasion fitness when it depends on multiple phenotypes that are influenced by the evolving locus. Specifically, we assume that the biological scenario is such that invasion fitness can be written as

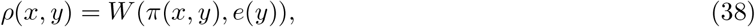

where

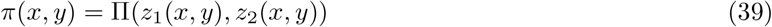

is the payoff to a heterozygote showing phenotypes *z*_1_(*x, y*) and *z*_2_(*x, y*), and *e*(*y*) is a regulating variable determined by the resident population (where individuals are homozygotes for the resident allele *y*). This form of invasion fitness, which is functionally equivalent to eq. (20), could for instance capture a situation where the fecundity of an individual depends on two of its phenotypes *z*_1_(*x, y*) and *z*_2_(*x, y*) (e.g. expression levels of two genes). Whatever the specific scenario, *z*_1_(*x, y*) and *z*_2_(*x, y*) are genetically correlated because they both depend on allelic values *x* and *y*. We assume that this dependency is given by

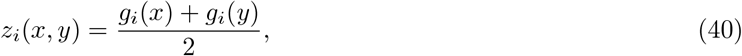

where *g*_*i*_(*h*) are the effects of an allele with genetic value *h* ∈ {*x, y*} on phenotype *i*. In other words, we assume additive genetic effects on the phenotype between homologous loci (which will allow us to connect to a number of existing results). These effects however may translate into non-additive effects on fitness.

Substituting eqs. (39)–(40) into eq. (21) shows that any singularity *y*\* must satisfy

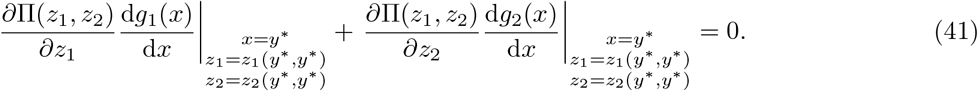

This says that at *y*\*, the effects on fitness of varying allelic value via phenotypes 1 and 2 must cancel each other out. Consider for example the case where fitness increases with both phenotypes (i.e. *∂*Π*/∂z*_1_ *>* 0 and *∂*Π*/∂z*_2_ *>* 0). Eq. (41) then tells us that for a singular trait value *y*\* to exist, phenotypes 1 and 2 need to change in opposite directions with *x*, namely 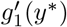 and 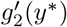 have opposite signs, such that a change in *x* has antagonistic effects on fitness via phenotypes 1 and 2: antagonistic effects which are balanced at *y*\*.

If we substitute eqs. (39)–(40) into eq. (22) and use eq. (41), we readily obtain that the interaction coefficient can be expressed as *I*_*π*_(*y*) = *L*_*π*_(*y*\*)*K*_*π*_(*y*\*)*/*2 where 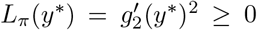, such that the sign of trait-dependent selection is governed by

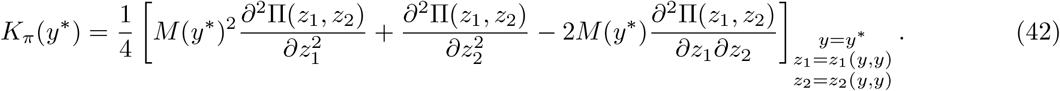

Here, *M* (*y*\*) = (*∂*Π*/∂z*_2_) */*(*∂*Π*/∂z*_1_)|_*y*=*y*_∗ gives the rate at which phenotype 1 can be substituted for phenotype 2 without changing payoff (i.e. the change in the number of units of phenotype 1 that correspond to the effect on payoff of a change in one unit of phenotype 2). Eq. (42) shows that NTDS tends to occur when: (i) the payoff function decelerates with phenotypes 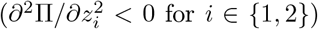, and/or (ii) phenotypes have complementary effects on payoff (*∂*^2^Π*/∂z*_1_*∂z*_2_ *>* 0). Both situations can lead to heterozygote advantage, i.e. where heterozygotes for divergent allelic effects close to the singular allelic value *y*\* have greater fitness than either homozygotes as a result of expressing an intermediate value for both phenotypes. In situation (i), when alleles contribute additively to two phenotypes (eq. 40), and payoff decelerates with both phenotypes in a concave way, heterozygotes can have greater fitness than either homozygotes (see Fig. 3 panel A for a graphical argument). In situation (ii), when the payoff exhibits complementary effects, this also favours a situation such that heterozygotes have a greater payoff (see Fig. 3 panel B for a graphical argument). Heterozygote advantage thus arises because the contribution of an allele to fitness is context-dependent, in the sense that its fitness effect varies across different phenotypes.

**Figure 3.**
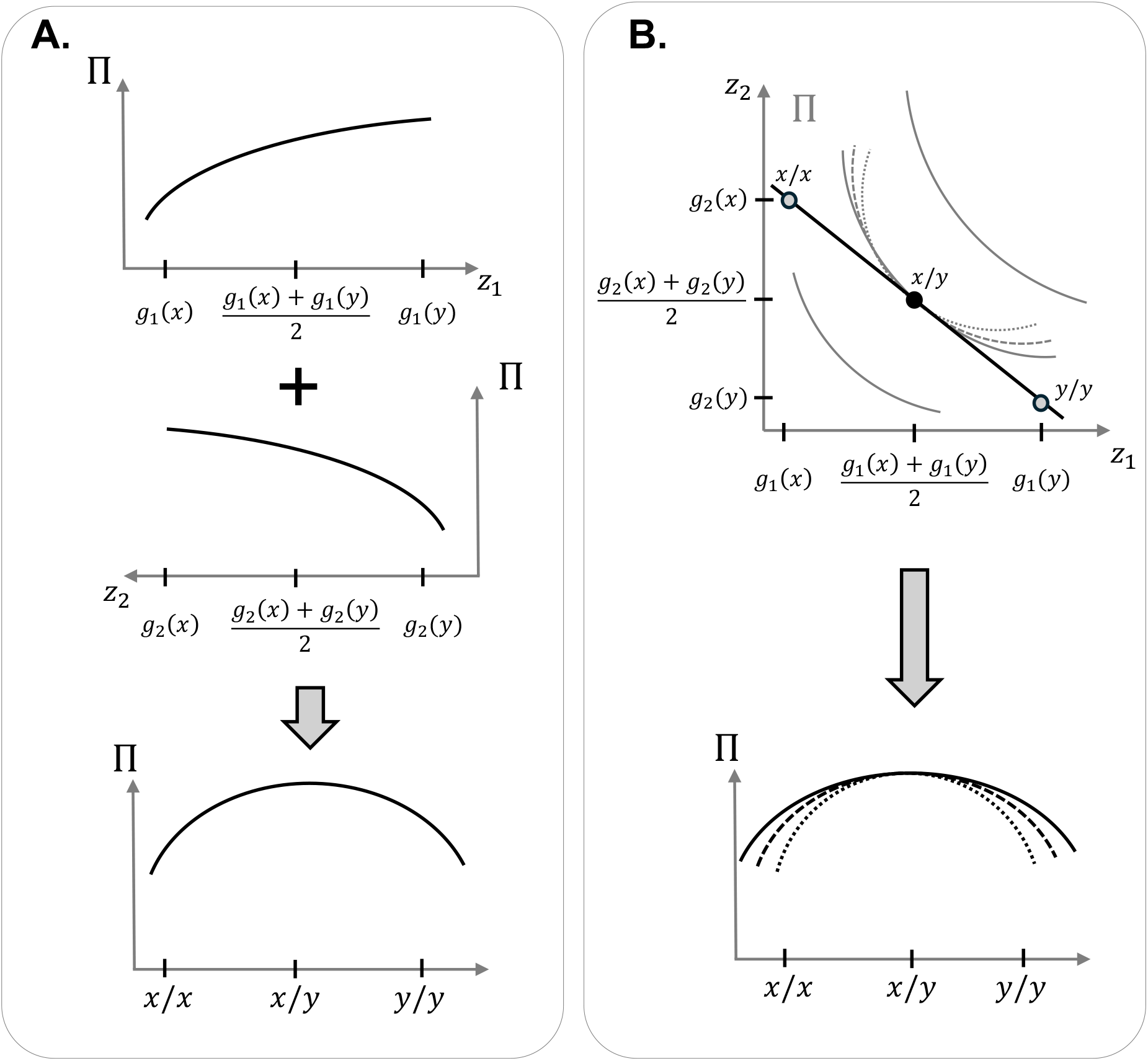
A graphical illustration of overdominance origin under pleiotropic constraints. **A.** Overdominance can arise when the payoff function Π(*z*_1_, *z*_2_) decelerates with both phenotypes increasing (first two terms of eq. 42). The top and middle graphs show the payoff Π(*z*_1_, *z*_2_) increasing in a decelerating manner with both *z*_1_ (top) and *z*_2_ (middle). When the payoff increases with both phenotypes, then *z*_1_ and *z*_2_ must change in opposite directions as a result of changing allelic value near a singular trait value (as implied by eq. 41). The direction of the x-axes are reversed in the top and middle graphs to ensure genotype alignment: *x/x* homozygotes on the left, *x/y* heterozygotes in the centre, and *y/y* homozygotes on the right. Summing the effects across both phenotypes to compute the total payoff for each genotype leads to overdominance, with heterozygotes (*x/y*) achieving the highest payoff (captured by the two first terms in eq. 42). **B**. Overdominance is also favoured when phenotypes have complementary effects on payoff (last term of eq. 42). The top graph shows a case where eq. (42) is negative and payoff increases with both phenotypes. Grey curves are payoff isoclines, which are convex owing to eq. (42) being negative (solid, dashed, dotted lines indicate increasing strength of complementarity). According to eq. (41), *z*_1_ and *z*_2_ vary in opposite directions near the singular trait value *y*\*, necessarily moving along tangents to a payoff contour (black lines, following the genotypes shown on graph). When mutant and resident allelic values *x* and *y* diverge in opposite directions away from *y*\*, this yields heterozygotes with higher payoff than either homozygote. The effect is stronger under complementarity effects (compare black line and grey curves in top panel and see bottom panel). This increase in mutant fitness under opposing allelic changes is a signature of NTDS.

This mechanism is in fact similar to the one described in the seminal SAS-CFF population genetic model (“Stochastic Additive Scale Concave Fitness Function”, Gillespie, 1978, 1991). That model assumes that alleles at a homologous locus contribute additively to physiological activity (which is a phenotype in our model, eq. 40) in a fashion that varies stochastically with environmental conditions (say phenotype 1 in environment 1 and phenotype 2 in environment 2); and that fitness increases in a decelerating–concave–manner with physiological activity. To see the connection with our approach here, consider that eq. (42) applies to this SAS-CFF scenario with Π taken as the geometric mean fecundity of the mutant and the regulating variable *e* as the geometric mean fecundity of a resident individual under a Wright-Fisher process (as assumed in Gillespie, 1978, 1991 see Appendix F.2 for details on how eq. 42 applies to the SAS-CFF scenario and how evolutionary branching can occur in this scenario).

While eq. (42) applies to a broader set of situations, its connection to the SAS-CFF model provides a couple of insights. Firstly, it reinforces the idea that heterozygote advantage should arise from context-dependent allelic effects that are summed over different contexts. In the original SAS-CFF model (Gillespie, 1978), these contexts are temporal (over changing environments), but the same principle applies across multiple phenotypes or biological functions. Secondly, since the SAS-CFF model is closely linked to the storage effect of ecological coexistence (Schreiber, 2020), this connection extends to our framework and can be phrased as follows. When conditions vary, diploidy allows alleles that are poorly adapted to current conditions to persist by buffering them in heterozygotes. This buffering effect can prevent the loss of alleles that may be beneficial under different environmental or functional conditions. In turn, the adaptive polymorphism that evolves under the above model leads to the divergence of two allelic values, each showing a better fit for these different environmental or functional conditions, showing heterozygote advantage in total.

### 6.2 Negative trait-dependent kin selection

We now consider the case where owing to limited dispersal, individuals interacting with one another show genetic associations generated by identity-by-descent, i.e., individuals are related (Hamilton, 1964; Michod, 1982 and see Rousset, 2004 for a review). This generalises the kin competition effects seen in sections 4.3-4.4. When relatedness is high, this means that a globally rare mutant is nevertheless likely to interact with other mutants locally. Relatedness should therefore modulate trait-dependent selection depending on the fitness effects of mutant-mutant interactions. To examine this, we analyse *I*(*y*\*) in the infinite island model of dispersal with two individuals per island (or patch or deme). Although the conditions for evolutionary branching in this model have been characterised previously (e.g. Day, 2001; Ajar, 2003), *I*(*y*\*) has not yet been analysed.

For simplicity and to focus on genetic interactions among individuals, we assume individuals are haploid and reproduce clonally. Invasion fitness in this case can be written as

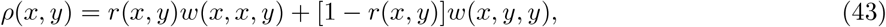

where *r*(*x, y*) is relatedness, that is, the probability that a rare mutant *x* in a resident *y* population has another mutant as patch neighbor (conversely, it has a resident as patch neighbor with probability 1−*r*(*x, y*)); *w*(*x, x, y*) is the individual fitness of a focal mutant *x* (first argument of *w*) when its patch neighbor is a mutant *x* (second argument), and the rest of the population in other patches consists of residents *y* (third argument); similarly, *w*(*x, y, y*) is the fitness of the mutant when its patch neighbor is a resident *y* (second argument).

Plugging eq. (43) into eq. (7a), we first recover the classical result that the singular trait value *y*\* must be such that the inclusive fitness effect, i.e. Hamilton’s rule in marginal form, vanishes:

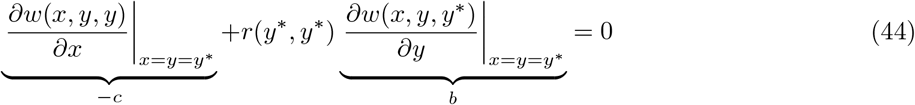

(e.g., Rousset, 2004). Therefore at *y*\*, the direct fitness effects of the trait (often conceived as a cost −*c*) must balance the relatedness-weighted indirect effects (i.e. the marginal effect a trait change in a neighbour has on an individual’s fitness; typically conceived as a benefit *b*), where relatedness here is evaluated in a population monomorphic for *y*\*. The equilibrium *y*\* can thus be seen as the result of a trade-off between different components of inclusive fitness: that is, between the effect an individual has on its own fitness and the effect it has on the fitness of relatives.

Substituting eq. (43) into eq. (6), we obtain that the interaction coefficient can be decomposed as

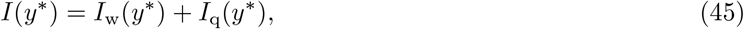

where

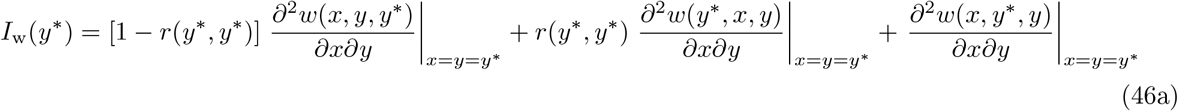

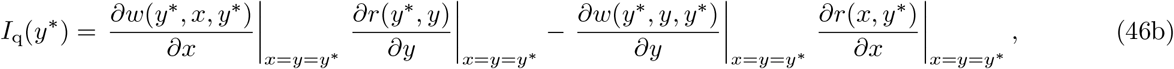

each capturing a pathway to negative trait-dependent selection.

The first term, *I*_w_(*y*\*), captures how invasion fitness changes when the mutant and resident traits are varied simultaneously in the same direction, while keeping relatedness constant, hence relatedness *r*(*y*\*, *y*\*) being evaluated in a population monomorphic for *y*\*. The first term of *I*_w_(*y*\*) shows NTDS is weakened in populations showing higher relatedness when joint trait changes in a mutant and its patch neighbour have negative fitness effects (i.e. where *∂*^2^*w*(*x, y, y*\*)*/∂x∂y <* 0; note that *∂*^2^*w*(*x, y*\*, *y*)*/∂x∂y*, in contrast, consists in the fitness effect of joint changes in the focal and individuals from *other* patches which are always non-relatives). Such a fitness effect could arise for various reasons. One relevant instance would be when the trait mediates a social interaction among patch neighbours that shows complementary or negative synergistic effects (as given by eq. 22, e.g. snowdrift games, Doebeli et al., 2004; Wakano and Lehmann, 2014, or trait based competition for limiting resources leading to character displacement, Ajar, 2003; Schmid et al., 2024). In that case, high relatedness makes polymorphism more difficult simply because it makes it more difficult for social partners to show differentiated phenotypes, and therefore enjoy the inherent benefit that being different yields under this type of interaction. Kin selection thus opposes NTDS here and this effect explains why several studies have found that limited dispersal tends to make evolutionary branching more difficult (e.g. Day, 2001; Ajar, 2003; Wakano and Lehmann, 2014; Mullon and Lehmann, 2019; Schmid et al., 2024). The next two terms of *I*_w_(*y*\*) consist of fitness effects of changes between individuals in different patches, which are expected to be weaker because interactions are indirect. Nevertheless, they indicate that high relatedness could in principle favour NTDS when the fitness effect of a neighbour’s trait depends negatively on trait expression in other patches (i.e. where *∂*^2^*w*(*y*\*, *x, y*)*/∂x∂y <* 0). To our knowledge, no existing models of polymorphism under limited dispersal invoke or seem to rely on this effect.

The second term of eq. (45), *I*_q_(*y*\*), reveals how kin selection can contribute to NTDS when relatedness depends on trait values. Suppose the trait has positive indirect fitness effects, so that increasing a neighbour’s trait value increases individual fitness (helping): *∂w*(*y*\*, *x, y*\*)*/∂x* = *∂w*(*y*\*, *y, y*\*)*/∂y >* 0 (evaluated at *x* = *y* = *y*\*). Then the first term of *I*_q_(*y*\*) shows that NTDS is promoted when increasing the population trait reduces relatedness among mutants, i.e. *∂r*(*y*\*, *y*)*/∂y <* 0. In that case, as the resident trait increases, mutants are less likely to interact with other mutants, so the indirect fitness benefit received from mutants showing increased helping is reduced. In other words, the gain from increasing the mutant’s trait declines as the resident trait increases, thus contributing to NTDS. Conversely, the second term of *I*_q_(*y*\*) shows that NTDS is promoted when increasing the mutant trait increases relatedness among mutants (i.e. *∂r*(*x, y*\*)*/∂x >* 0, e.g. because higher mutant trait values increase philopatry). In that case, increasing the resident trait increases the help a mutant receives conditional on interacting with a resident neighbour, but this effect is weaker when the mutant trait is higher because higher *x* increases relatedness and therefore reduces the probability that a mutant interacts with resident neighbours.

The term *I*_q_(*y*\*) shows that relatedness can either promote or oppose NTDS, depending on how trait expression feeds back on inclusive fitness. This relies on traits affecting both the fitness of neighbours and the probability that those neighbours are relatives. An example of such a process leading to the evolution of a social–dispersal syndrome is given in Appendix F.3. More generally, any trait that modifies relatedness–such as those affecting budding dispersal, dispersal success, establishment probability, generational overlap, or group extinction–may generate negative trait-dependent kin selection.

## 7 The role of modifiers of genetic context in NTDS

We saw in section 6.2 how group structure and the heterogeneity it entails allow for a form of NTDS that is relatedness-mediated. There, it arose when trait expression increased individual fitness in one context (e.g. when being in a group with individuals expressing a similar trait due to identity-by-descent) and the population trait moving in the same direction reduces the probability of individuals being in that context (e.g. due to dispersal decreasing identity-by-descent). This mechanism rests on the existence of heterogeneity in the population: mutant individuals do not interact randomly but instead experience different fitness effects depending on the local genetic composition of their group or patch. More generally, population heterogeneity and the existence of different contexts in which individuals can find themselves open new pathways for NTDS: it can arise whenever a trait influences both the contexts an individual experiences and fitness within those contexts.

Box 3 formalises this idea by considering populations structured into discrete classes to capture heterogeneity either within or between generations. These classes may correspond to different genetic backgrounds, phenotypic states, environmental conditions, or stages in an individual’s life cycle. The general form of the interaction coefficient in such populations (eqs. 3.E–3.F) shows that trait-dependent selection is influenced not only by how population trait expression affects fitness within each class but also how it modifies the class structure. In particular, NTDS emerges when an increase in the resident trait value reduces either: (i) the probability that mutants occur in the class in which their fitness is increased owing to a mutant trait increase; or (ii) the reproductive value of individuals produced by mutants in that class (see Box 3 for further details).

The mechanism we have described here should be especially relevant to the maintenance of polymorphism in traits that influence how individuals experience environmental or demographic variation, such as sex ratio, habitat choice, bet-hedging mechanisms, aging, or life-history switches. Consider for instance a trait influencing habitat choice and reproductive success in that habitat, such that mutants expressing greater trait value preferentially settle in a specific habitat that confers them higher fitness (Ravigné et al., 2009). However, if the population also expresses higher trait values and preferentially settles in the same habitat, this increases local competition and thus reduces the per capita benefit of the trait, leading to NTDS. More broadly, these considerations show how population heterogeneity should favor diversity by allowing different morphs to specialise in different contexts, favouring the coexistence of different trait values that associate their reproductive success with the conditions where they perform best [see also Avila and Mullon, 2023 for similar considerations of context-dependence for the coefficient *H*(*y*\*)].

## 8 Discussion

Polymorphism is widespread in nature. Numerous models from population genetics, evolutionary ecology, and game theory have proposed mechanisms by which polymorphisms can be maintained by natural selection. Here, we have aimed to organise these mechanisms by the common thread of negative trait-dependent selection (NTDS), which helps to clarify when adaptive polymorphism can be expected under the gradual evolution of quantitative traits.

### 8.1 Monomorphic instability, the role of NTDS, and its underlying mechanisms

We started by noting that characterizing the adaptive polymorphism favoured by long-term evolution is, in general, a computationally hard problem (section 2.2.3). This difficulty perhaps explains why the bulk of theoretical models about quantitative trait evolution focus on directional selection or on identifying singular trait values in monomorphic populations. Indeed, the phenotypic selection gradient is the central tool to understand adaptation, from evolutionary ecology (Vincent and Brown, 2005; Pásztor et al., 2016), through kin selection (Frank, 1998; Rousset, 2004), to quantitative genetics (Lande, 1980; Arnold, 2023). Nevertheless, necessary conditions for the emergence of polymorphism by gradual evolution can be derived from the evolutionary instability of convergence stable singularities (section 2.3). This requires the selection interaction coefficient to be negative, which, biologically, means that higher population trait values reduce the marginal fitness gain from further increases in the trait. Equivalently, opposite changes in mutant and resident trait values contribute positively to invasion fitness through their interaction. This negative trait-dependent selection (NTDS) is a necessary condition for adaptive polymorphism (eq. 8). As such, NTDS provides a general computable condition to identify candidate polymorphisms under gradual evolution, independent of specific biological mechanisms.

To clarify the biological origins of NTDS, we structured invasion fitness around two classes of fitness factors (section 3). Production factors are inputs to an individual’s survival and reproduction that depend directly on its own trait (e.g. baseline fecundity, germination rate). Regulating factors, meanwhile, are inputs to survival and reproduction that depend on the traits expressed by conspecifics (e.g. local density, other’s behaviour). These two types of fitness factors can act at different biological levels: from phenotypes, to payoffs from interactions, to invasion fitness (Fig. 1).

When regulating factors at the level of invasion fitness depend solely on the population trait (e.g. eq. 10), our model captures ecologically mediated interactions. We showed that NTDS here requires an interaction between trait-induced trade off in production and trait-driven constraints in regulating factors (eq. 14). This enables niche partitioning, with different trait values adapting to different environments associated with different regulating variables. Focusing on NTDS allowed us to uncover a previously undescribed life-history polymorphism under a standard fecundity–survival trade-off (e.g. Charnov, 1993; Case, 2000). Limited dispersal, by generating local kin competition, enables the coexistence of a morph with low survival and high fecundity alongside a quasi-immortal morph with low fecundity when external mortality is very low. The robustness of this deserves further investigation.

Our general analysis allows for regulating factors depending on population and mutant traits (section 3.3), unlike earlier related formalisations of invasion fitness (e.g. Metz, 2011; Pásztor et al., 2016; Lion, 2018). This generalisation is useful for capturing several biologically relevant scenarios: (1) behavioural responsiveness, where resident individuals adjust their actions in response to mutant traits; (2) diploid inheritance, where mutant alleles co-occur with resident alleles in heterozygotes; and (3) mutant–mutant interactions, as occur under limited dispersal, family interactions, or assortative mating. In such cases, NTDS can arise through interaction effects on regulating factors, without requiring direct effects on production factors. We illustrated these broader mechanisms through examples involving socially and genetically mediated interactions.

Socially mediated interactions involve direct behavioural encounters between individuals (e.g. Maynard Smith, 1982; Akçay and Van Cleve, 2012; McNamara and Leimar, 2020). When fitness increases with the payoff obtained from such interactions, and regulating factors at the fitness level depend only on the trait of the resident population (due to e.g. density-dependent regulation), the interaction coefficient can be evaluated directly at the level of payoffs (eq. 22). However, the regulating variables that determine these payoffs may themselves depend on mutant traits. This occurs, for instance, when individuals modulate their behaviour in response to their partner’s traits, i.e. under social responsiveness. Social responsiveness introduces additional pathways to NTDS, which can act independently of direct trait effects on payoffs (eq. 27). This led us to revisit models of the evolution of traits that bias how individuals evaluate the consequences of their own actions in pairwise interactions under complete information (section 5.3.2). These dispositions influence not only an individual’s own behaviour but also, via social responsiveness, the behaviour of their partners. When interactions have antagonistic effects on payoff (as per eq. 31), we found that selection can favour a dimorphism between “optimists,” who overvalue the payoff benefits of their actions, and “pessimists,” who undervalue them.

Finally, we considered how genetically mediated interactions can influence NTDS. One relevant instance of this is interactions between mutant and resident alleles within diploid individuals. Under random mating, conditions for the emergence of polymorphism depend solely on heterozygote fitness, making them equivalent to conditions identified in haploid or asexual systems (eq. 36). This implies that diploidy, in itself, does not change the ecological or social conditions under which gradual evolution leads to polymorphism. Therefore, if one’s interest lies primarily in understanding adaptive polymorphism, explicit modelling of genetic architecture is unnecessary: a form of phenotypic gambit for adaptive trait variation. Nevertheless, diploidy can generate NTDS at the level of alleles close to a singularity when there are pleiotropic constraints over multiple production factors (e.g. survival and fecundity). In particular, such constraints can lead to overdominance, or heterozygote advantage, a classical mechanism maintaining genetic variation. Our analysis clarified that, under gradual evolution, complementary allelic effects within heterozygotes (rather than simply intermediate optimal phenotypes) are required for overdominance to drive diversification (eq. 42).

Another important type of genetic interaction arises when limited dispersal generates identity-by-descent between interacting individuals so that kin selection occurs (e.g. Hamilton, 1964; Michod, 1982; Rousset, 2004). Here, we revealed a form of trait-dependent kin selection, which when negative, can contribute to diversification. Specifically, our analysis demonstrates that NTDS can emerge when a trait simultaneously increases the fitness of neighbouring individuals and local relatedness (eq. 46). Such traits effectively alter their own genetic context, creating conditions in which indirect fitness benefits are disproportionately high when the mutant and resident are different. More broadly, when population structure creates different contexts in which the trait can be expressed and influence fitness (e.g. sex, age or environmental structure), NTDS is increased if the trait expressed by an individual increases its reproductive success in a particular context and trait expression at the population level reduces the representation of individuals in this context (Box 3). This pathway, which appears to be underappreciated in the context of NTDS, should be particularly relevant for traits influencing habitat choice, senescence, and sex determination, as these traits naturally interface with context-dependent selection. In this sense, population heterogeneity creates distinct niches, allowing genetic effects to specialise within particular contexts, a process through which diversity begets diversity.

### 8.2 Oligomorphic stability and open questions

Once selection favours the emergence of polymorphism via evolutionary branching (i.e. when *H*(*y*\*)+*I*(*y*\*) *<* 0 and *H*(*y*\*) *>* 0), an essential question is where the evolutionary dynamics will lead the population in the long-run. Comparatively few results are available to characterize these long-term outcomes of adaptive polymorphism in quantitative traits. Nevertheless, a useful insight is that such a polymorphism should be oligomorphic so that only a discrete number of morphs stably coexist in the uninvadable coalition (e.g. Ellner and Sasaki, 1996; Gyllenberg and Meszéna, 2005). For most of our illustrative examples, we analyzed this oligomorphism in the Appendix (Appendix D.1 and Appendix E.1). But in general, analytically characterizing this oligomorphism, especially when environments fluctuate through time, remains challenging. A relevant research avenue would therefore be developing approaches or techniques to better infer and characterize the convergence stable and uninvadable coalition.

For different examples analysed in the Appendix, we found that at the convergence stable and uninvadable coalition, there is no trait-dependent selection in the sense that *I* = 0 at evolutionary equilibrium (see Appendix E.2.1 and Appendix E.2.2). This pattern has been established previously in scenarios where regulating factors depend solely on resident trait values and where the number of coexisting morphs matches the number of independent regulating variables (Kisdi and Geritz, 2016, Proposition 6). In such cases, the resulting polymorphism is said to be saturated (Kisdi and Geritz, 2016). Our analysis suggests that saturation–in the broader sense of all trait-dependent selection vanishing–can occur under more general conditions. Future work could therefore aim to establish sufficient conditions under which adaptive evolutionary endpoints result in vanishing trait-dependent selection, going beyond ecologically mediated interactions. Such knowledge would be particularly valuable because it would allow evolutionary end-points to be fully characterized by the selection gradient alone (since local uninvadability coincides precisely with convergence stability under vanishing trait-dependent selection). Given that the most biologically explicit representation of the selection gradient is the generalized marginal inclusive fitness effect for polymorphic populations (which is capable of capturing both class and group structure; Priklopil and Lehmann, 2021, eq. 101-102), this would further solidify the central role that Hamilton’s marginal rule plays in understanding the nature and process of adaptation of quantitative traits.

### 8.3 Conclusions

By decomposing invasion fitness into production and regulating factors and focusing on negative trait-dependent selection (NTDS) we obtained necessary conditions for adaptive polymorphism applicable across diverse biological systems. This allows for (i) a consistent criterion to identify the presence or absence of NTDS; (ii) a systematic way of categorising regulating factors at the phenotype, payoff, and fitness levels, clarifying their roles in driving NTDS; and (iii) a synthesis of ecological, social, and genetic scenarios that are conducive to NTDS and thus adaptive polymorphism. These illustrative scenarios, while not exhaustive, highlight the diverse origins of polymorphism. They reinforce the notion that given the hierarchical organisation of biological systems and the ubiquity of competition, NTDS may be common rather than exceptional. If NTDS is indeed so prevalent, it may hold the key to understanding why intra-specific trait diversity is so pervasive, continuously providing the fuel for evolutionary change.

## Acknowledgements

We thank Claus Rueffler for many useful discussions, providing references, and spotting an oddity within a previous version of one model. We gratefully acknowledge two anonymous reviewers who provided extremely useful and super extensive comments that fundamentally improved the manuscript, the results, and the presentation.

## Appendix A Uninvadable coalition existence

In Box 1, we state that when 𝒳 is non-empty and compact and the invasion fitness *ρ*(*x, p*) is continuous in *x* and *p* [meaning the function *ρ*: 𝒳 × Δ(𝒳) → ℝ_+_ where Δ(𝒳) is equipped with the weak* convergence topology, Alipantris and Border, 2006, 15.1, is jointly continuous], then a mixed Nash equilibrium satisfying eq. (5) is guaranteed to exist. This follows from applying established results to the best-reply correspondence 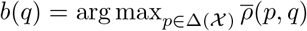 as is usually done in game theory for normal form games (Fudenberg and Tirole, 1991; Mas-Colell et al., 1995). Here, 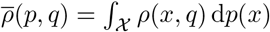 and see eq. (A-8) for a concrete representation of the invasion fitness *ρ*(*x, q*).

When 𝒳 is non-empty and compact, so will be Δ(𝒳) (Alipantris and Border, 2006, Th. 15.11). By Berge’s maximum theorem (Alipantris and Border, 2006, Th. 17.31), the best-reply *b*(*q*) then defines an upper hemi-continuous and non-empty compact-valued correspondence. Because *b*(*q*) is convex-valued (since 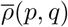 is linear in *p*) and Δ(𝒳) is a convex set (since the mixture of two probability distributions is a probability distribution), the conditions of Glicksberg’s fixed point theorem are satisfied, and therefore the set of equilibrium points satisfying *p*\* ∈ *b*(*p*\*) is non-empty and compact (Alipantris and Border, 2006, Th. 17.55).

In biological terms, there is therefore at least one uninvadable coalition in any evolutionary model that satisfies our main biological assumptions (section 2) in the absence of temporal environmental fluctuations (adaptive polymorphism characterized by eq. 3), provided that 𝒳 is compact and *ρ*(*x, p*) is continuous in *x* and *p*. This uninvadable coalition may consist of an adaptive polymorphism or be a singleton trait, say *x*\*, in which case the uninvadable trait value satisfies *x*\* ∈ arg max_*x*∈𝒳_ *ρ*(*x, x*\*) for invasion fitness *ρ*(*x, y*) of a mutant *x* in a monomorphic resident population *y*. This latter case corresponds to a symmetric Nash equilibrium in pure strategies and such an equilibrium exists if *ρ*(*x, y*) is continuous, quasi-concave in its first argument, and 𝒳 is compact and convex (e.g. Fudenberg and Tirole, 1991, p. 34). It would be useful to extend these existence results to the case of temporally fluctuating environments (adaptive polymorphism characterized more generally by eq. 2).

## Appendix B Dimorphic coexistence near a singularity

Under gradual evolution, evolutionary convergence towards an invadable monomorphism *y*\* results in the population subsequently splitting into two distinct morphs leading to their coexistence in a protected polymorphism (Eshel et al., 1997; Geritz et al., 1998). This coexistence can be understood by considering two morphs, say *x*_1_ = *y* + *δ* ∈ 𝒳 and *x*_2_ = *y* − *δϵ* ∈ 𝒳, that are weakly diverged around *y* with |*δ*| ≪ 1, where *ϵ >* 0 is assumed to be of order *O*(1). These morphs will coexist if they can mutually invade one another, that is when *ρ*(*x*_1_, *x*_2_) *>* 1 and *ρ*(*x*_2_, *x*_1_) *>* 1 (e.g. Hofbauer and Sigmund, 1998; Turelli et al., 2001). Second order Taylor expanding *ρ*(*x*_1_, *x*_2_) and *ρ*(*x*_2_, *x*_1_) in *δ* around *δ* = 0, and using the condition *ρ*(*y, y*) = 1 when differentiating twice with respect to *y*, we get

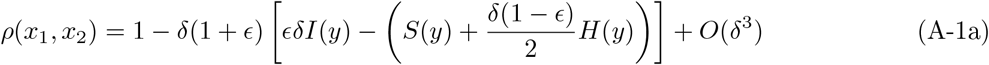

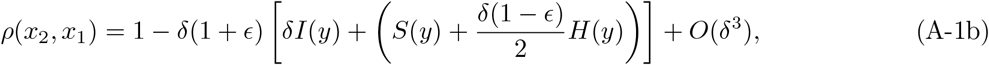

where *S*(*y*), *H*(*y*) and *I*(*y*) are given in eq. (6). The term (*S*(*y*) + *δ*(1 − *ϵ*)*H*(*y*)*/*2) has opposite effects on the invasion fitness of each morph: increasing fitness of *x*_2_ and diminishing that of *x*_1_. Any mechanism that makes this term closer to zero can therefore be seen as an equalizing mechanism in the jargon of coexistence theory (Chesson, 2000b). Equalizing mechanisms reduce inherent fitness differences among morphs thus favouring coexistence. From that perspective, directional selection leading to a singular trait value *y*\* such that *S*(*y*\*) = 0 can be seen as an equalizing mechanism.

The term *I*(*y*), in contrast, influences the invasion fitness of each morph in the same way. Any mechanism that makes *I*(*y*) more negative thus acts as a stabilizing mechanism in the jargon of coexistence theory, favouring mutual invasion. When *I*(*y*) *>* 0, mutual invasion is not possible, i.e. *I*(*y*) *<* 0 is a necessary condition for mutual invasion. Additionally, when *ϵ* = 0 (such that the two morphs are symmetrically distributed around *y*) and *y* = *y*\* is singular, then *I*(*y*) *<* 0 also becomes a sufficient condition for mutual invasion.

## Appendix C Oligomorphic coexistence

We here extend the analysis of the (in)stability of monomorphism (section 2.3) to the case of oligomorphism. This is to (i) complement the analysis of the main text, (ii) show how the decomposition of invasion fitness eq. (9) can be applied to the oligomorphic case, (iii) allow us to characterize the uninvadable polymorphism for examples in the main text, and (iv) display some of the complexities involved in characterizing an uninvadable and convergence stable oligomorphism.

### Appendix C.1 No class structure nor environmental fluctuations

We start by describing the conditions for the uninvadability and convergence stability of an oligomorphism for the case of absence of (i) class structure and (ii) between generation fluctuating environments (and discuss relaxing these assumptions in Appendix C.2). For this case, we can replace *µ* by *p* in eq. (2) and since we consider only a resident population with a discrete number *n* of segregating morphs we set *p* = **p** = (*p*_1_, *p*_2_, …, *p*_*n*_) to denote the probability distribution of these morphs in the population 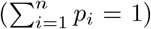, with support 𝒫(**p**) = **y** = (*y*_1_, *y*_2_, …, *y*_*n*_). We then denote by 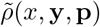 the invasion fitness of type *x* in a type **y** population with morph frequency distribution **p**.

#### Appendix C.1.1 Uninvadable oligomorphism

At an equilibrium of the dynamics of morph frequencies, each morph *y*_*i*_ ∈ 𝒫(**p**) must have invasion fitness equal to one so that the equilibrium frequency vector **p** must satisfy:

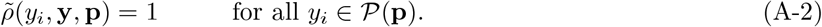

Assume such an equilibrium exists and that it is hyperbolic (a hyperbolic equilibrium means that the Jacobian matrix of the maps 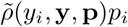 of the dynamical system of morph frequencies *i* = 1, .., *n* has eigenvalues with non-zero real parts, Hirsch et al., 2004). Hyperbolicity guarantees that (i) the equilibrium **p**(**y**) of eq. (A-2) can be written as a continuous function of **y** and (ii) that any of the finite number of fixed point of eq. (A-2) retain their stability properties upon changes of morph **y** values around a singularity **y**\* (from the implicit function theorem Sydsaeter et al., 2008, p.84 and Principle 3 of Karlin and McGregor, 1972). Then, the invasion fitness of a mutant *x* ∈ 𝒳 can be written as

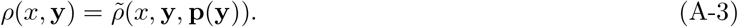

From this, the three coefficients of selection of eq. (6) generalize to

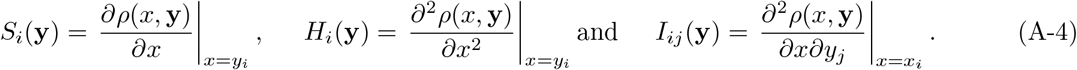

A sufficient condition for an oligomorphic resident population with support 𝒫(**p**\*) = **y**\* and equilibrium frequency **p**\* to be locally uninvadable with trait values in the interior of the state space is thus that for all *i* ∈ {1, 2, …, *n*}:

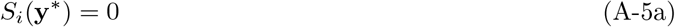

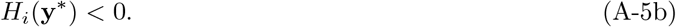

A necessary condition for local uninvadability is that eq. (A-5b) holds with weak inequality. An oligomorphism satisfying only condition (A-5a) is referred to as a singular coalition, and those satisfying both conditions (A-5a) and (A-5b) as locally uninvadable coalitions. The concept of a locally uninvadable coalition is standard, e.g. eq. (17) in Vincent et al. (1996), eq. (15) in Geritz et al. (1998), eq. (36) in Sasaki and Dieckmann (2011), or eqs. (11) in Kisdi and Geritz (2016).

#### Appendix C.1.2 Convergence stable oligomorphism and saturation

When conditions (A-5a) and (A-5b) hold, a coalition with morphs **y**\* and frequencies **p**\*, if established, will no longer change. But will such a pair (**y**\*, **p**\*) establish through gradual evolution; namely, is (**y**\*, **p**\*) an attractor of the evolutionary dynamics? Convergence stability here means that for any coalition **y** ≠ **y**\* in a small neighborhood of **y**\*, the evolutionary dynamics should gradually converge to (**y**\*, **p**\*) by small step mutations. In a population of morphs **y** ≠ **y**\* within some neighborhood of the uninvadable coalition, the associated equilibrium morph frequencies **p** ≠ **p**\* will differ from that of the uninvadable coalition. If **y**\* is favoured in such a population, the frequency of morphs should eventually be attracted to **p**\*, which requires that this is a local asymptotically stable fixed point of morph frequency dynamics. We therefore require that an uninvadable coalition **y**\* is locally convergence stable in the sense that it is a phenotypic attractor and that **p**\* is a local asymptotically stable point of morph frequency dynamics (implying that if any of the regulating variables are extensive variables, then they must also reside at asymptotically stable fixed points of their local dynamics).

In a population at **y** close to an uninvadable **y**\* with local asymptotically stable **p**\*, the selection gradient *S*_*i*_(**y**) is sufficient to describe the direction of long-term evolution. Specifically, *S*_*i*_(**y**) determines whether a mutant *y*_*i*_ + *δ* arising from morph *y*_*i*_ with |*δ*| ≪ 1 in population state **y** will go extinct or substitute the ancestral type, leading to a new polymorphic population state (*y*_1_, *y*_*i*−1_, *y*_*i*_ + *δ, y*_*i*+1_, …, *y*_*n*_) (Cai and Geritz, 2020; Priklopil and Lehmann, 2020, 2021). Since mutations occur independently in each morph, the rate of trait change of morph *i* is given by *v*_*i*_(**y**(*t*))*S*_*i*_(**y**), where *v*_*i*_(**y**(*t*)) *>* 0 is some measure of genetic variance in the subpopulation of morph *i*. More often than not, the variances *v*_*i*_(**y**(*t*)) differ among different morphs (due to e.g. different subpopulation sizes of morphs or selection, Sasaki and Dieckmann, 2011). Therefore, a necessary condition for a polymorphic singularity satisfying eq. (A-5a) to be convergence stable, i.e. to be attracted towards **y**\* regardless of the scaling factor, is that the Jacobian matrix

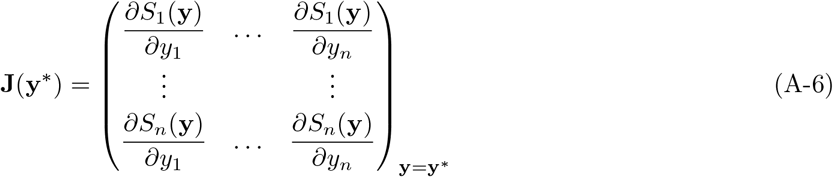

has all its eigenvalues with negative real parts. Sufficient conditions for convergence stability include (i) that **J**(**y**\*) is a negative definite matrix (i.e., all eigenvalues of the symmetric part of 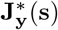 are negative in which case the oligomorphic singularity is said to be strongly convergence stable (Apaloo and Butler, 2009; Kisdi and Geritz, 2016) and (ii) that **J**(**y**\*) is a sign stable matrix (e.g. Quirk and Ruppert, 1965, Theorem 5, May, 1973, p. 71), in which case convergence stability can be predicated from qualitative properties of the signs of the entries of 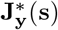.

In the same way as eq. (7b) decomposes convergence stability into quadratic selection and trait-dependent selection, the Jacobian eq. (A-6) can be decomposed as

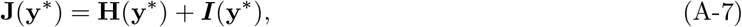

where **H**(**y**\*) is a diagonal matrix with (*i, i*)-element *H*_*i*_(**y**\*) capturing disruptive selection on morphs *i* (eq. A-4), while matrix ***I***(**y**\*) has (*i, j*)-element given by *I*_*ij*_(**y**\*), which is the effect of a change in the trait of morph *j* on selection in morph *i*. As such, matrix ***I***(**y**\*) captures trait-dependent selection in an oligomorphic population. For an oligomorphism **y**\* to be locally convergence stable yet invadable (so that *H*_*i*_(**y**\*) *>* 0 for some *i*), it is necessary that ***I***(**y**\*) contains negative entries and thus that some NTDS occurs. To see this, suppose that all non-diagonal elements of ***I***(**y**\*) are non-negative. In this case, **J**(**y**\*) is a Metzler matrix. We then know that for **J**(**y**\*) to have all eigenvalues with negative real parts, it is necessary that **J**(**y**\*) has only negative diagonal elements (Briat, 2017, Lemma 3.1). This entails that it is necessary that *I*_*ii*_ *<* 0 for any trait *i* where *H*_*i*_(**y**\*) *>* 0. This latter condition also holds if each morph evolves independently of each other (Geritz et al., 1998; Sasaki and Dieckmann, 2011).

#### Appendix C.1.3 Model in terms of fitness factors and vanishing NTDS

The expression of invasion fitness in terms of production and regulating factors for a monomorphic population (eq. 9) immediately extends to the oligomorphic case by setting

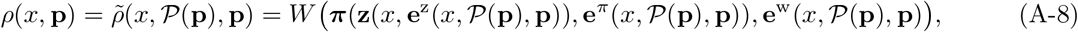

where the regulating factors are now expressed in terms of the set of traits **y** = 𝒫(**p**) in the population and their frequency distribution **p**. This representation of invasion fitness also applies more generally to the case where there is a continuum number of traits 𝒳 ⊆ ℝ in the population with distribution **p** = *p*, as originally introduced in section 2.2.1. We thus note that in Box 1, where we mention continuity of *ρ*(*x, p*) with respect to (*x, p*) for existence of Nash equilibria, this then means continuity of 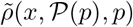 with respect to *p* it the third argument in a concrete model.

For the case of ecologically mediated interaction (section 4) invasion fitness reduces to 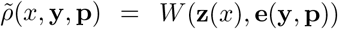 and the population trait feedbacks only act through the regulating variables. Interestingly, Kisdi and Geritz (2016, Proposition 6) have shown that for this case and when the number of oligomorphs in the uninvadable coalition matches the number of regulating variables, then trait-dependent selection vanishes; that is

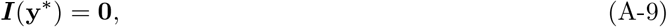

where **0** is a matrix of zeroes. Then, the criteria for convergence stability collapses to that of local uninvadability (recall eq. A-7), and in the terminology of Kisdi and Geritz (2016), the resulting adaptive polymorphism is said to be saturated.

### Appendix C.2 Class structure and environmental fluctuations

In the presence of class structure (*sensu* Box 3), but still no environmental fluctuations, the broad line of argument described in Appendix C.1.1–Appendix C.1.2 still applies, but then eq. (A-2) needs to be amended by adding an auxiliary system of equations to describe the equilibrium allele frequency in each class (see Lion et al., 2022; Bukkuri and Brown, 2023 for work on the stability of oligomorphism in class structured populations).

In the presence of between generation environmental fluctuations, there will no longer be an evolutionary equilibrium in morph values **y**\* associated with some deterministic morph frequency equilibrium **p**\*. This frequency has to be replaced by some stationary ergodic probability distribution 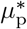 over morph frequencies. In principle, the selection coefficients defined in eq. (A-4) can be applied to characterize directional selection, uninvadability, trait-dependent selection, and convergence stability under environmental fluctuations (by extending eq.A-2 to environmental fluctuations). However, we are aware of no example where this has been achieved analytically without further approximations (see Ellner and Sasaki, 1996 for work on a concrete characterization of the stability of an oligomorphism under between generation environmental fluctuations and Ellner, 1996 for extending this to class structured population).

## Appendix D D Ecologically mediated interactions

### Appendix D.1 Interaction coefficient

We derive here eqs. (13)–(14) of the main text. We first substitute eq. (10) into eq. (6), which after factoring out common terms gives

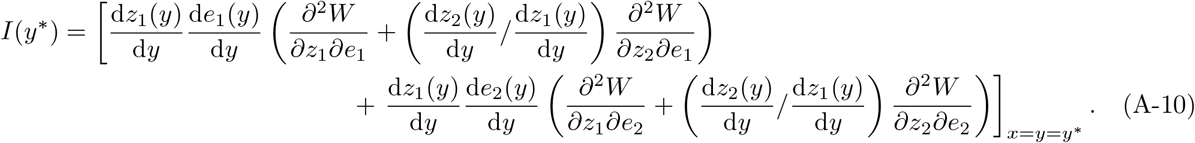

Then, using eqs. (11)–(12) to substitute 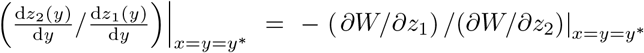 and 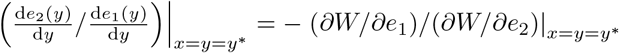 produces the desired result by noting that

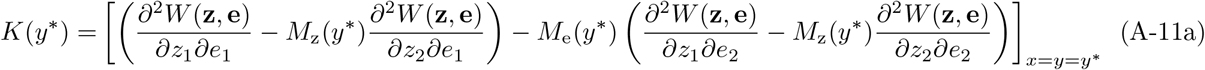

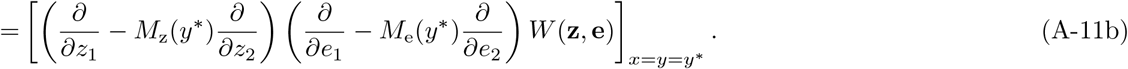

### Appendix D.2 Life-history evolution and limited dispersal

Substituting eq. (16) into eq. (11) shows that any singularity *y*\* must solve

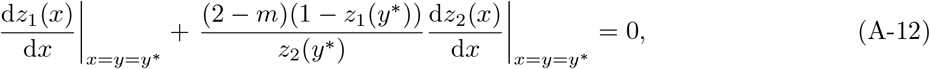

which expresses the trade-off between investing in survival (first term) and reproduction (second term). The smaller (2 − *m*), the larger the investment in fecundity required to compensate for a given loss of survival. Since (2 − *m*) is smallest when dispersal is unlimited (*m* = 1), individuals in this case evolve to invest less in reproduction than under limited dispersal (all else equal). As explicit fitness factors, we use

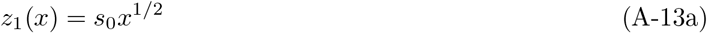

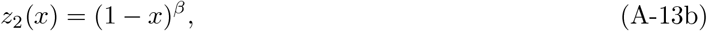

with 0 *< β* ≤ 1 and 0 *< s*_0_ ≤ 1. Note that if *s*_0_ = 1, then putting all resources into survival means the individual becomes immortal, and this can be thought of as a situation where there is no external source of mortality. We analyse the model with *s*_0_ close to one. Using eq. (A-13) in eq. (A-12) reveals a unique feasible singular level of reproductive effort,which can be written as

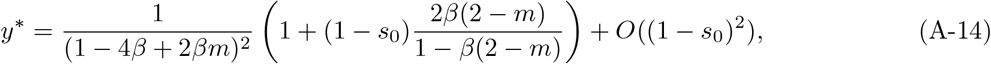

where the remainder is a complicated function of the parameters. Using eq. (6) produces

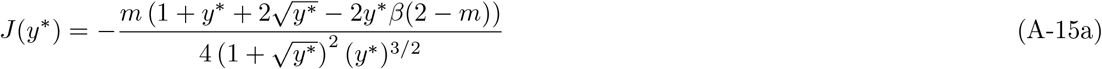

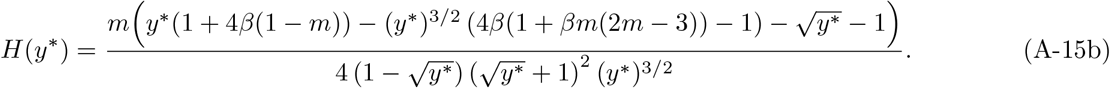

It can be checked that there is a range of parameter values in *m* and *β* such that *J*(*y*\*) *<* 0 and *H*(*y*\*) *>* 0, whereby dimorphism is favored by gradual evolution at *y*\* and this can occur only when *s*_0_ is close to one. For instance when *m* = 0.6 and *s*_0_ = 0.996, polymorphism is favoured as long as 0.664 *< β <* 0.755, and when *β* = 0.7, the singular trait value is *y*\* = 0.8264 from which evolutionary branching will occur (see Fig. 2 for corresponding individual-based simulations).

Given branching occurs, let us now consider evolution in a polymorphic population with trait values 𝒫(**p**) = **y** = (*y*_1_, *y*_2_, …, *y*_*n*_) (as described in Appendix C). Generalizing eq. (16) to the polymorphic case, the invasion fitness of type *x* in a polymorphic population is then given by

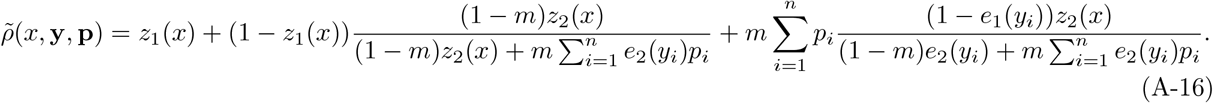

Because there are only two regulating factors in this model, let us focus on a dimorphism, *n* = 2, whereby using *p*_2_ = (1 − *p*_1_) and applying eq. (A-2) to eq. (A-16) with eq. (A-13) shows that

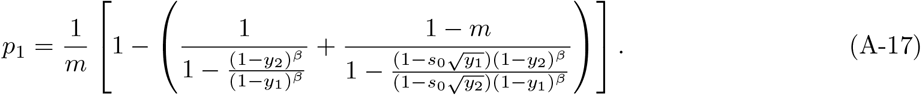

Evaluating the selection gradients (*S*_1_(*y*_1_, *y*_2_), *S*_2_(*y*_1_, *y*_2_)) on morphs *y*_1_ and *y*_2_ (by applying eq. (A-4) on eq. (A-16) with *n* = 2 together with eq. (A-13) and eq. (A-17)), we were not able to find analytical expressions for the singular coalition 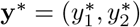. But numerical analysis allows to characterize it, and for the parameter values *s*_0_ = 0.996, *m* = 0.6 and *β* = 0.7, we find a unique (symmetric) singular coalition 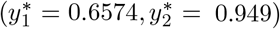 with frequency 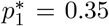 of morph one, consistent with corresponding individual-based simulations (Fig. 2). While polymorphism obtains in this model only over a small range of parameter values, these results suggests that adaptive polymorphism of life-history traits under limited dispersal merits further investigations.

We finally observe that, in the special case where *s*_0_ = 1, starting from a convergence stable singular point, selection pushes morph 2 towards the boundary of the trait space at 1, such that this morph tends to become immortal. Meanwhile, morph 1 evolves towards a mixed strategy with an ever-decreasing frequency and eventually vanishes (using L’Hôpital’s rule in eq. (A-17) under *s*_0_ = 1 and 1 *> β >* 0, we have 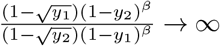 as *y*_2_ → 1, and therefore *p*_1_ → 0 as *y*_2_ → 1). It can then be checked that selection on morph 1 itself vanishes as *y*_2_ → 1 (*S*_1_(*y*_1_, *y*_2_) → 0 as *y*_2_ → 1). However, mutations should preclude convergence to *y*_2_ = 1 exactly. Hence, the coexistence of a quasi-immortal morph and a second morph investing in a more balanced way in both survival and reproduction appears possible.

### Appendix D.3 Temporal heterogeneity and limited dispersal

Substituting fitness model eq. (18) with 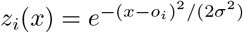 into eqs. (6)–(7a) shows that there is a unique singular strategy

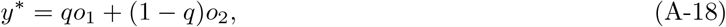

and that

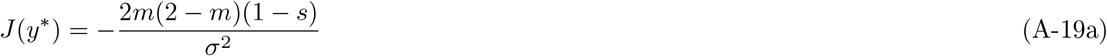

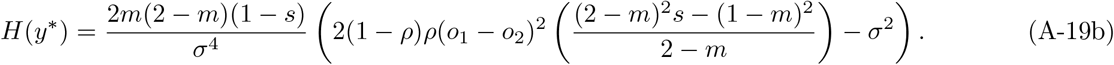

This shows that *y*\* is convergence stable for *s* ∈ [0, 1) and that evolutionary branching can occur under a certain range of parameter values. The larger *s* is, the more likely polymorphism is to emerge, while a decrease in *m* inhibits branching. We do not attempt to characterize the resulting oligomorphism here, because the presence of environmental fluctuations make this problem challenging (see Ellner and Sasaki, 1996 for the case when *m* = 1).

## Appendix E Socially mediated interactions

### Appendix E.1 Payoff based gradients

We here derive condition (22). For this, first substitute eq. (20) into the selection gradient of eq. (6) and use the chain rule to see that it can be written in terms of the payoff gradient as

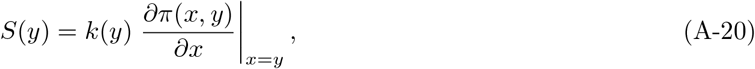

where *k*(*y*) = *∂W* (*π, e*(*y*))*/∂π*|_*π*=*π*(*y,y*)_ *>* 0 by assumption. Then, any singular strategy *y*\* satisfying *S*(*y*\*) satisfies

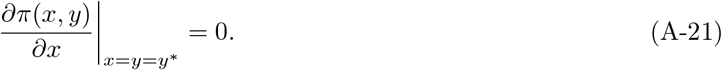

Using this equality after having substituted eq. (20) into the second-order selection coefficients of eqs. (6)–(7a), we find that

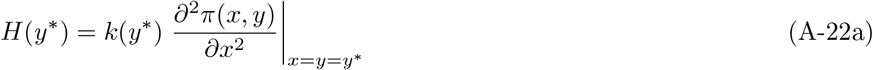

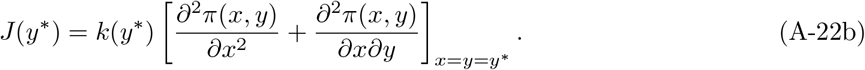

Hence, the Hessian, interaction and Jacobian coefficients can all directly be computed from the payoff function. The necessary condition *I*(*y*\*) = *k*(*y*\*)*I*_*π*_(*y*) *<* 0 for protected polymorphism can then be reduced to ineq. (22), since *k*(*y*\*) *>* 0.

### Appendix E.2 Examples: contests and preference evolution

#### Appendix E.2.1 Contest evolution

Substituting payoff model eq. (28) into eqs. (A-21)-(A-22) and using *a*(*x*) = *e*^*ωx*^ along *C*(*a*(*x*)) = *x*^2^, we get a unique singular point

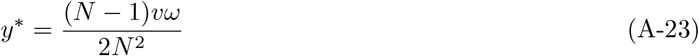

and

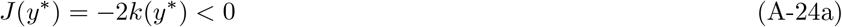

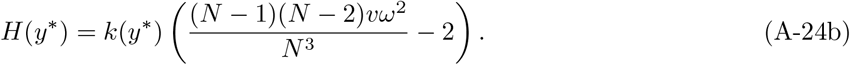

Thus, when *N* ≤ 2, the singular trait value is always convergence stable and uninvadable. When *N >* 2, one can find parameter values such that *H*(*y*\*) *>* 0. Thus, it is clear from the second line of eq. (A-24) that there is a range of parameter values where a dimorphism can be favored by gradual evolution at *y*\*. This is for instance the case when *N* = 5, *ω* = 2, and *v* = 10, which is confirmed by individual-based simulations in Fig. 2. Analysing the resulting dimorphism mathematically, however, would be entirely numerical, as the invasion fitness of type *x* in a polymorphic population relies on multinomial sampling of the different morphs to determine the payoff of individuals during contests. We do not pursue this approach here, as the next example can be solved entirely analytically and thus illustrates how to evaluate asymptotic polymorphism under socially mediated interactions.

#### Appendix E.2.2 Preference evolution: dispositions

Here, we analyze the model of dispositions of section 5.3.2 but extend the payoff of this model to

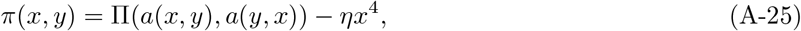

where *a*(*x, y*) is given by eq. (33) and the term *ηx*^4^ represents a cost of overconfidence, which increases with *x* and is controlled by the parameter *η* ≥ 0. This payoff implements the main text model eqs. (30)–(31) but adds a cost to dispositions. This prevents the open ended evolution of exaggerated trait expression. The right hand side of eq. (A-25) is therefore more general than the right hand side of eq. (23). As a result the first-order condition for model eq. (A-25) is not given directly by eq. (24). Nevertheless the synergy coefficient corresponding to eq. (A-25) is equivalent to that obtained from model eqs. (30)–(31).

Substituting payoff model eq. (A-25) along equilibrium actions (33) into eq. (21) yields a unique feasible singular strategy *y*\*, which can be expressed as

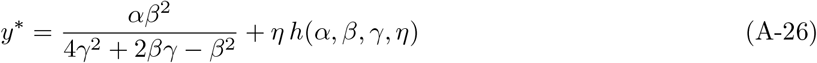

for some complicated function *h*(*α, β, γ, η*) of the parameters. When *η* = 0 and *γ* = 1, *y*\* reduces to the level of disposition found by Heifetz et al. (2007a, p. 267). The Jacobian and the Hessian coefficients at this singular point are obtained from eq. (A-22) to read as

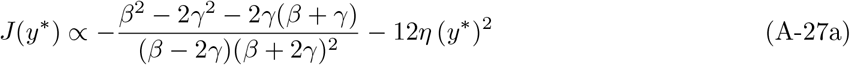

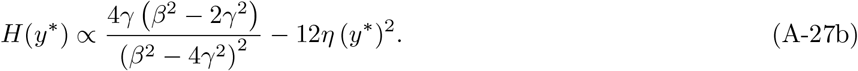

Straightforward analysis of these equations reveals that there is a range of parameter values such that *J*(*y*\*) *<* 0 and *H*(*y*\*) *>* 0 and so evolutionary branching will occur. For instance, this is the case when *α* = 0.2, *β* = 1.6, *γ* = 1, and *η* = 0.45.

Given that branching occurs, let us consider evolution in a polymorphic population 𝒫(**p**) = **y** = (*y*_1_, *y*_2_, …, *y*_*n*_). Assuming a Wright-Fisher process with fecundity *f*_b_ + *π*(*x, y*) of an individual with trait *x* interacting with one with trait *y* (where *f*_b_ ≥ 0 is some baseline fecundity allowing for payoff to be negative), the invasion fitness of type *x* in a polymorphic population is using eq. (A-25) along eq. (31) given by

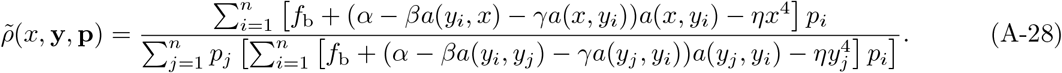

Focusing on a dimorphism, *n* = 2, and solving 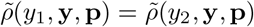 for *p*_1_ by setting *p*_2_ = 1 − *p*_1_, we find a unique complicated analytical expression for *p*_1_(*y*_1_, *y*_2_). Nevertheless, this expression allows us to compute the selection gradients *S*_1_(**y**) and *S*_2_(**y**) analytically. By solving *S*_1_(**y**\*) = 0 and *S*_2_(**y**\*) = 0 for 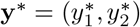, we find a single singular coalition characterized by

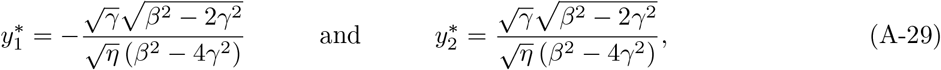

which leads to

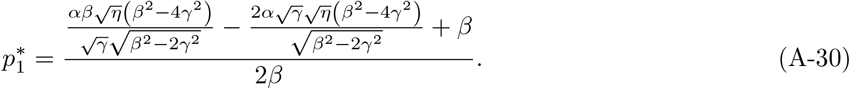

Computing the Hessian for each morph (eq. 7a), we find that

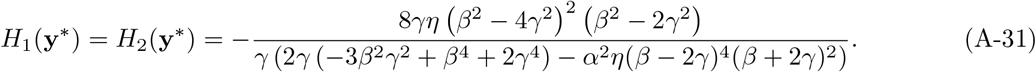

Computing the Jacobian matrix **J**(**y**\*) (eq. A-7) shows that its two eigenvalues take the same value as the Hessian coefficients eq. (A-31). Hence, whenever the coalition is locally uninvadable, it is also strongly convergence stable and the polymorphism is saturated, i.e. eq. (A-9) is satisfied. This is the case for instance with the aforementioned parameter values (*α* = 0.2, *β* = 1.6, *γ* = 1, and *η* = 0.45), which leads to a symmetric uninvadable coalitions with trait values 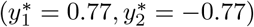, which is confirmed by individual-based simulations in a population of size *N* = 10000 with baseline fecundity *f*_b_ = 4 (see Fig. 2).

#### Appendix E.2.3 Preference evolution: other-regarding motivations

Here, we consider an alternative model of preference evolution, in which the evolving trait *x* measures an individual’s regard for others. We assume that the utility functions for a pair of interacting individuals, one mutant with action *a* and trait *x* and a resident with action *b* and trait *y*, are given by

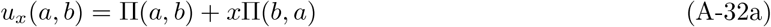

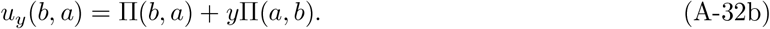

Each utility is the sum of own material payoff to the actor plus the payoff to the interaction partner and thus represents an other-regarding preference. The weight of other-regard is the evolving traits. To close the model, we postulate the linear-quadratic payoff eq. (31) of the main text. The necessary first-order conditions for a Nash equilibrium under this game are that *∂u*_*x*_(*a, b*)*/∂a* = 0 and *∂u*_*y*_(*b, a*)*/∂b* = 0, which can be expressed as

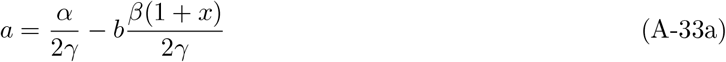

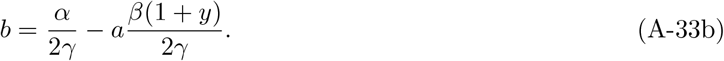

Solving for the equilibrium actions *a* = *a*(*x, y*) and *b* = *a*(*y, x*) yields

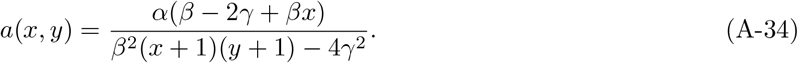

This is indeed a Nash equilibrium action since the solution is unique and satisfies the second-order condition *∂*^2^*u*_*x*_(*a, b*)*/∂a*^2^ = −2*γ* ≤ 0 and *∂*^2^*u*_*y*_(*b, a*)*/∂b*^2^ = −2*γ* ≤ 0.

For this model, the corresponding singular strategy, response coefficient and action interaction coefficients are

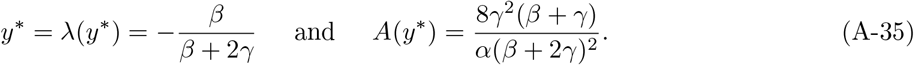

Since *β >* 0 and *γ >* 0, the singular strategy is always convergence stable and *∂*^2^Π(*a, b*)*/*(*∂a∂b*) *<* 0, *∂*Π(*a, b*)*/*(*∂b*) *<* 0, [*∂*^2^Π(*a, b*)*/∂a*^2^ + *∂*^2^Π(*a, b*)*/∂b*^2^] = −2*γ <* 0, and *∂*Π*/∂a* + *∂*Π*/∂b* = −*αβ/*2*γ <* 0. This entails that the two components of eq. (26) result in negative trait-dependent selection:

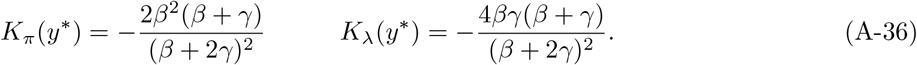

Polymorphism in other-regard can thus evolve and it can be checked that there is a range of parameter values where evolutionary branching occurs, leading to the open-ended evolution of a positive *x >* 0 and negative *x <* 0 other-regarding morph unless a cost is added to the expression of the trait.

**Appendix F Genetically mediated interactions**

### Appendix F.1 No effect of diploidy on the instability of monomorphism

We here show eq. (36). For this, first substitute eq. (35) into the selection gradient of eq. (6) and use the chain rule to show that it can be written as

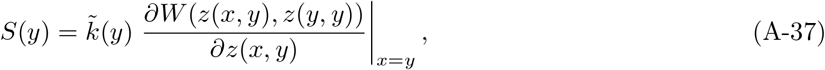

where 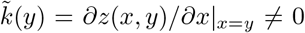 by assumption. Thus, any singular strategy *y*\* satisfying *S*(*y*\*) = 0 has to satisfy

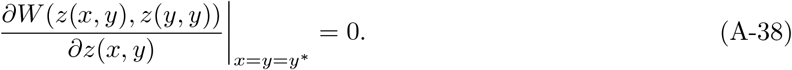

Using this equality along *∂z*(*x, y*)*/∂x*|_*x*=*y*_ = *∂z*(*x, y*)*/∂y*|_*x*=*y*_ by substituting eq. (35) into the second-order coefficients of eqs. (6)–(7a), we find that

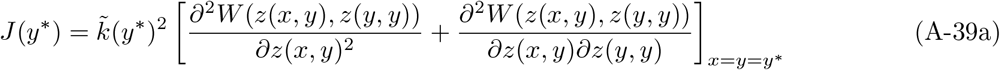

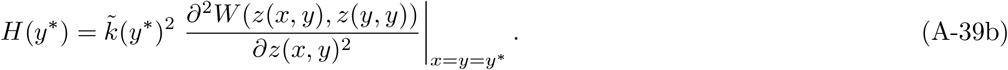

Hence, for *J*(*y*\*) *<* 0 and *H*(*y*\*) *>* 0 to hold, requires that

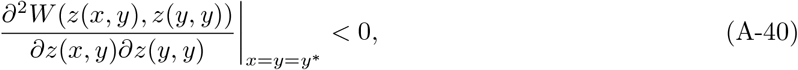

i.e. there is NTDS at the phenotypic level.

Because in eqs. (A-38)–(A-40) all derivatives are phenotypic and these phenotypes are evaluated at *z*(*y*\*, *y*\*) = *z*\*, the allelic value *y*\* is in fact not needed to perform the local evolutionary analysis, which can be carried out by focusing only on the phenotype value *z*\* solving

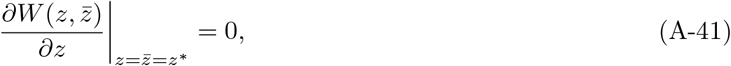

and this leads to condition (36) of the main text.

The above shows that under our assumptions, diploidy does not affect the conditions for the gradual emergence of polymorphism for evolutionary branching. However, diploidy does affect the conditions for mutual invasibility of alleles *x* and *y*. From eq. (A-1) the mutual invasibility of two weakly and symmetrically diverged allelic values around *y*\* requires that

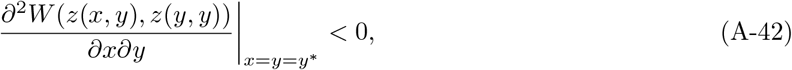

which holds if and only if

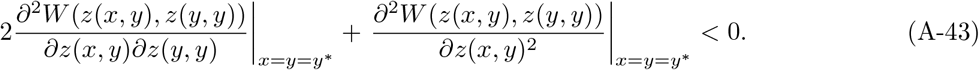

This, in particular, captures the notion that with diploidy, two alleles on either side of a local fitness peak (i.e., either side of a singular value *y*\* such that *∂*^2^*W* (*z*(*x, y*), *z*(*y, y*))*/*(*∂z*(*x, y*)*∂z*(*y, y*)) = 0 and *∂*^2^*W* (*z*(*x, y*), *z*(*y, y*))*/∂z*(*x, y*)^2^ *<* 0 at *x* = *y* = *y*\*) can be maintained owing to overdominance: i.e. where the intermediate heterozygote expresses a fitter phenotype. However, such polymorphism is necessarily shortlived on an evolutionary timescale as any allele that codes for the optimal phenotype in homozygous form will invade and fix.

### Appendix F.2 Average heterozygote advantage

We here detail an example leading to heterozygote advantage through temporal averaging over different contexts by considering a scenario envisaged by Gillespie (1978), where the environment changes each generation, being in state 1 with probability *q* and in state 2 with probability 1 − *q* such that invasion fitness is given by

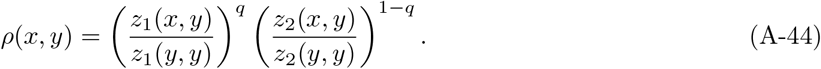

This takes the form of eq. (38) and (39) with 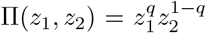 and 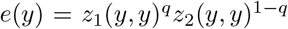 under the assumption of a Wright-Fisher process: *ρ*(*x, y*) = *π*(*x, y*)*/π*(*y, y*) where *π*(*y, y*) = *e*(*y*). The interpretation is that *z*_1_ and *z*_2_ are the physiological activity of a gene in environments 1 and 2, respectively; and we assume that activity increases with allelic value in environment 1, say 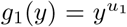, and decreases with allelic value in environment 2, say 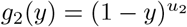 (with 0 *< y <* 1).

Using 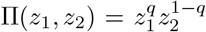 in eq. (39) when substituting eq. (40) along 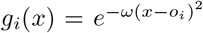 and eq. (38) into eqs. (6)–(7a) shows that there is a unique singular strategy

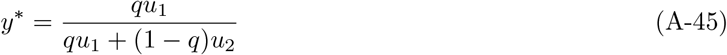

and that

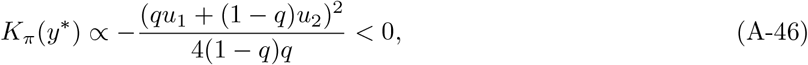

which displays NTDS. Further,

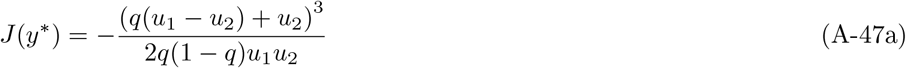

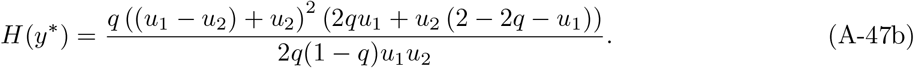

Thereby, provided *u*_*i*_ *>* 2 and *u*_*j*_*/*(2*u*_*i*_) *>* (1 − *q*)*/*(*u*_*i*_ − 2*q*) (for *i* ≠ *j*), evolutionary branching will occur and the coexistence of two morphs: one that increases activity in environment 1 and another in environment 2, leading to heterozygote advantage on average over the two environments. See also Siljestam and Rueffler (2024) for a conceptually similar model that leads to poly-allelic polymorphism when there are more than two biological functions of a gene.

### Appendix F.3 Trait-dependent kin selection: social-dispersal syndrome

As an example of relatedness modulated NTDS, let us consider a situation where the trait constitutes a form of helping that increases the fecundity of one’s neighbour, and that its costs are over dispersal such that an individual who invests more resources to help one’s neighbour has fewer resources to commit to its dispersing offspring. Trait expression therefore has positive indirect fitness benefits (via fecundity) and positive effects on relatedness (via dispersal), especially when rare. For simplicity, we assume a Moran type life cycle whereby exactly one individual dies per life-cycle iteration in each patch, such that individual fitness takes the form

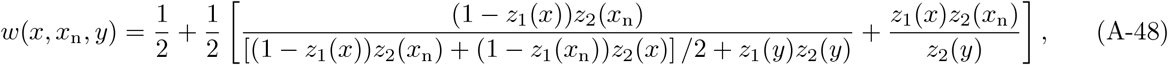

where *z*_1_(*x*) is the probability of dispersal of an individual expressing trait value *x* (which we assume is a decreasing function) and *z*_2_(*x*_n_) is the fecundity of an individual whose neighbour expresses trait *x*_n_ (an increasing function of *x*_n_). Therefore, the first term within square bracket in eq. (A-48) is the philopatric component of fitness, i.e. the expected number of offspring that establish in the natal patch, and the second term is the dispersal component, i.e. the expected number of offspring that establish in other patches. Using Box 2 of Alger and Lehmann, 2023, we obtain that relatedness for this model can be expressed as

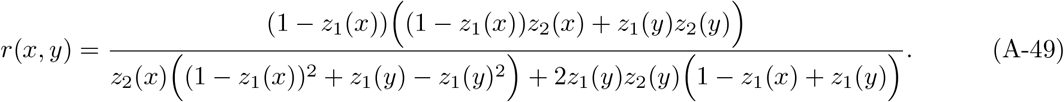

Strictly speaking, this is a relatedness proxy, whereby the right-hand side of eq. (43) becomes the basic reproductive number of the mutant, which can be used to predict evolutionary outcomes, since the basic reproductive number is sign equivalent to invasion fitness, e.g., Metz et al. (2008) and see Mullon et al. (2016) for these connections under limited dispersal.

Analysis of this model does not give rise to a simple expression for the interaction coefficient *I*(*y*\*) from which useful information can be extracted, but a straightforward numerical approach with *z*_1_(*x*) = 1 − *x* and 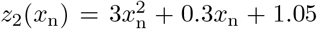 shows that at *y*\* = 0.38, we have *I*(*y*\*) *<* 0 while *I*_q_(*y*\*) *<* 0 *< I*_w_(*y*\*), i.e. the population experiences NTDS due to relatedness effects only. Additionally, the population undergoes evolutionary branching leading to the coexistence of two differentiated morphs that show a social-dispersal syndrome: (i) one that helps and thus tends to remain philopatric (expressing *x* ≈ 0.65); and (ii) another that does not help and disperses (expressing *x* = 0). Trait-dependent selection can be understood by considering the mutant and resident population both being more selfish and thus more prone to dispersal simultaneously. Such a mutant benefits by dispersing away and thereby avoiding interactions with other mutants. However, if the resident population simultaneously becomes more selfish and dispersal-prone, these benefits vanish because resident individuals help less anyway and also disperse. In that case, both mutant and resident lineages experience the same reduction in relatedness, so the mutant does not differ in its ability to interact preferentially with relatives. A handful of models have reported the evolution of similar social-dispersal syndromes owing to kin selection (Mullon et al., 2018; Prigent and Mullon, 2023) but beyond these, there remain few analyses of how effects on relatedness can generate NTDS.

## Appendix G Class structured populations

### Appendix G.1 Within generation heterogeneity

Here, we show eq. (3.E) in Box 3. To that end, we first note that eq. (3.A) of that box can be written as

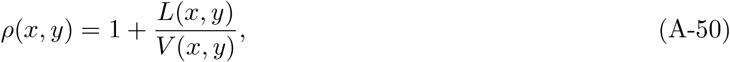

with

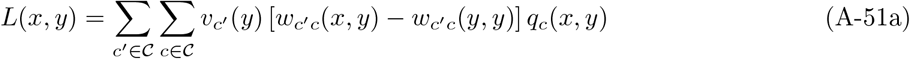

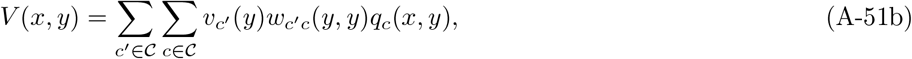

(see Ohtsuki et al., 2020 for a proof), where *v*_*c*_(*y*) is the (neutral) reproductive value of an individual in class *c* in a resident population *y* defined to satisfy the recursion *v*_*c*_(*y*) = ∑ _*c*_′_∈𝒞_ *v*_*c*_′ (*y*)*w*_*c*_′_*c*_(*y, y*) and assumed to satisfy the normalization ∑ _*c*∈𝒞_ *v*_*c*_(*y*)*q*_*c*_(*y, y*)=1, where *q*_*c*_(*y, y*) satisfies *q*_*c*_′ (*y, y*) = ∑ _*c* ∈𝒞_ *w*_*c*_′_*c*_(*y, y*)*q*_*c*_(*y, y*) (e.g., Taylor, 1990; Rousset, 2004). For completeness, note that one also has *q*_*c*_′ (*x, y*)*ρ*(*x, y*) = ∑ _*c*∈𝒞_ *w*_*c*_′_*c*_(*x, y*)*q*_*c*_(*x, y*), whereby *ρ*(*x, y*) is the leading eigenvalue of the matrix with elements *w*_*c*_′_*c*_(*x, y*) that has leading right eigen-vector with elements *q*_*c*_(*x, y*) [and summing the equation over all *c*^′^ ∈ 𝒞 produces eq. 3.A, see Lehmann et al., 2016; Ohtsuki et al., 2020 for more details on invasion fitness derivation and the interpretations of the weights *q*_*c*_(*x, y*)].

The choice of scaling fitness effects with reproductive values satisfying the aforementioned normalization ensures that under neutrality (*x* = *y*), one immediately has *V* (*y, y*) = 1 and note that the term in squared brackets in eq. A-51a is zero and thus *L*(*y, y*) = 0 at *x* = *y*. Using these equalities after differentiating *ρ*(*x, y*) as given by eq. (A-50) with respect to both *x* and *y*, we obtain,

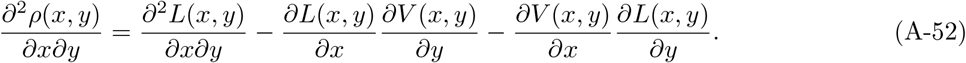

Now note that *S*(*y*) = *∂L*(*x, y*)*/∂x* and that by total differentiating *ρ*(*y, y*) with respect to *y* one obtains that *∂L*(*x, y*)*/∂x* = −*∂L*(*x, y*)*/∂y*. Since at a singular point *S*(*y*\*) = 0, we have

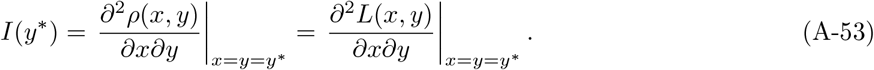

Using eq. (A-51a) for *L*(*x, y*) we thus obtain

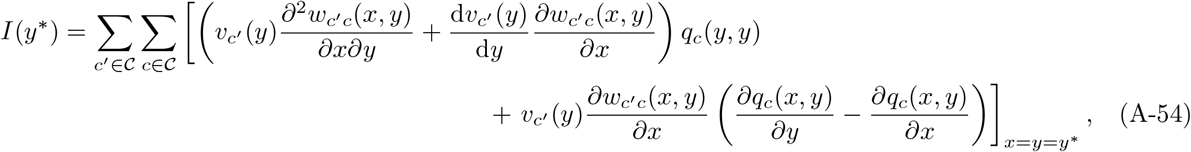

where the last term comes from differentiating [*w*_*c*_′_*c*_(*x, y*) − *w*_*c*_′_*c*_(*y, y*)] *q*_*c*_(*x, y*) in eq. (A-51a) with respect to *x* and *y* but considering only derivatives involving perturbations of the class frequency state *q*_*c*_(*x, y*) (thus excluding *∂*^2^*w*_*c*_′_*c*_(*x, y*)*/∂x∂y*). The term [*w*_*c*_′_*c*_(*x, y*) − *w*_*c*_′_*c*_(*y, y*)] reflects the fact that in a population showing within generation heterogeneity, natural selection depends on class specific fitness differences (e.g. Priklopil and Lehmann, 2024, p. 225), which translates in perturbations of class frequency states contributing to the interaction coefficient depending on differences in marginal effects on class frequency states, i.e., the *∂q*_*c*_(*x, y*)*/∂y* − *∂q*_*c*_(*x, y*)*/∂x* term in eq. (A-51a). Finally, by factoring out *q*_*c*_(*y, y*) and *v*_*c*_′ (*y*) in eq. (A-51a) and using the property of the logarithm function, we obtain eq. (3.E) of the main text.

### Appendix G.2 Between generation heterogeneity

We here show eq. (3.D) and eq. (3.F) of Box 3. For this, it is useful to express invasion fitness as *ρ*(*x, y*) = *e*^*ϕ*(*x,y*)^ where *ϕ*(*x, y*) is the invasion exponent of the mutant *x* in a resident *y* population (e.g. Tuljapurkar, 1990; Ferrière and Gatto, 1995). Owing to the fact that *ϕ*(*y, y*) = 0, we obtain by direct differentiation

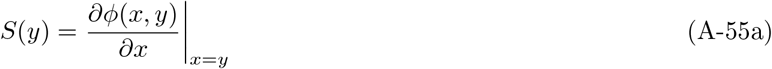

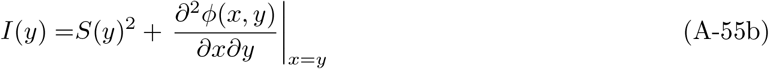

and at a singularity *I*(*y*\*) = *∂*^2^*ϕ*(*x, y*)*/∂x∂y*| _*x*=*y*=*y*_* since *S*(*y*\*) = 0. Thus, we can compute all derivatives of invasion fitness directly from all corresponding derivatives of the growth rate, which facilitates calculations in what follows.

Substituting eq. (3.B) into *ϕ*(*x, y*) = log(*ρ*(*x, y*)), we get upon rearrangement

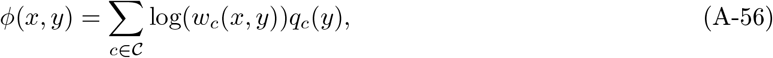

which is the familiar expression for the mutant growth rate under between generation heterogeneity (e.g. Haccou and Iwasa, 1995, eq. 6). Differentiating, we get

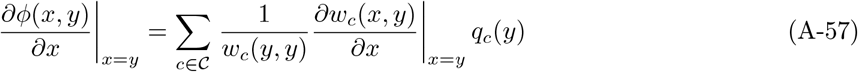

and

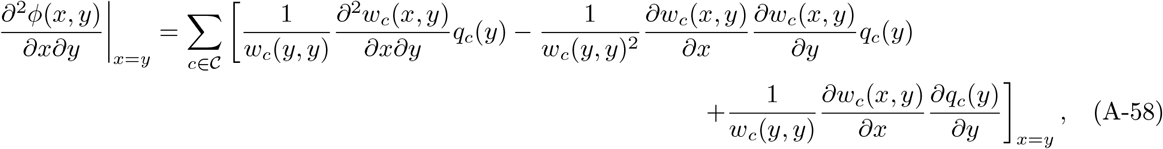

which, after re-arrangements by factoring out *q*_*c*_(*y*) and using the property of logarithm gives eq. (3.F).

### Appendix G.3 Class-specific regulation and production factors

Following the same construction of fitness factors as in section 3.2, we can also express *w*_*c*_′_,*c*_(*x, y*) and *q*_*c*_(*x, y*) in terms of fitness factors as follows by writing

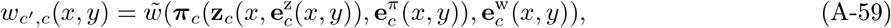

where we have indexed by *c* the production and regulating factors by the class of the bearer of this fitness, since in class structured population they will be class specific as both the phenotypes expressed by an individual and its experienced environment may depend on the class in which the individual resides (but should in principle not depend on the offspring class). The right hand side of eq. (A-59) can also be applied to fitness *w*_*c*_(*x, y*) for temporal heterogeneity.

The distribution *q*_*c*_(*x, y*) represents the likelihood of being in a certain state. Since the occurrence of any state can depend on all factors affecting reproduction and survival across all states, we generally have

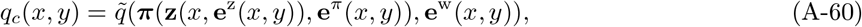

for some function 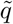 mapping the collection of all fitness factors to probabilities of state occurrence, and where a vector **v** without subscript (where **v** could be ***π*, z, e**^z^, **e**^*π*^, or **e**^w^) concatenates the vectors with subscript, i.e. **v** = {**v**_*c*_}_*c*∈𝒞_. By substituting eqs. (A-59)–(A-60) into eq. (3.A), we can express invasion fitness as in eq. (9), i.e., as a mapping from all fitness factors to geometric growth ratio (and the same logic applies to *w*_*c*_(*x, y*) and *q*_*c*_(*y*) under temporal heterogeneity).

Box 1. Uninvadable coalition, mean invasion fitness, Nash equilibrium, and computational complexity

The correspondence between uninvadability of a coalition (eq. (3)) and mean invasion fitness maximization (eq. (5)) extends to the case of between generation environmental fluctuations (i.e., when the uninvadable coalition is given by eq. (2)). To work this out, let ℳ = Δ(Δ(𝒳)) denote the set of probability distributions (measures) over the set Δ(𝒳) of population distributions. That is, *µ*_p_ ∈ ℳ is a random probability measure. Then, the mean fitness of measure *v* ∈ ℳ in a population with measure *µ*_p_ ∈ ℳ is given by

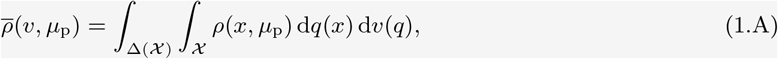

where *q* ∈ Δ(𝒳). With this, an uninvadable distribution *µ*\*_p_ satisfies eq. (2) if and only if it satisfies

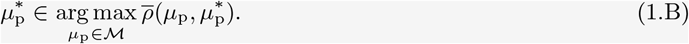

In other words, 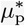 is an uninvadable distribution if and only if it maximizes mean invasion fitness in a population with distribution 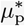, whereby 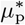 is a best response to itself. This correspondence between eq. (2) and eq. (1.B) can be understood by applying the standard relationship between pure and mixed strategy Nash equilibria from game theory [Mas-Colell et al., 1995, p. 250 and Narahari, 2014, p 99 for discrete strategy (trait) space, and for metric spaces, see Anderson et al., 2026, Proposition 1]. Namely, eq. (2) says that all morphs (pure strategies) in the set of segregating morphs 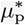 induced by 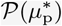 are uninvadable under population distribution 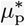 (a mixed strategy). To see that this is equivalent to the distribution *µ*\*_p_ being a best response to itself in terms of mean invasion fitness consider that eq. (1.B) holds but that 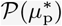 contains a morph that is invadable under 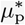 and thus has invasion fitness lower than 1 (i.e. eq. (2) does not hold). Then, the average population invasion fitness could be increased by decreasing the frequency of this morph and concomitantly increasing the frequency of any morph that is uninvadable. This then contradicts the assumption that the population distribution 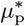 is a best response to self [and thus eq. (1.B) implies eq. (2)]. Conversely, if all morphs in 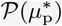 are uninvadable under 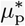 (eq. (2) holds), then the population distribution 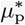 smust be a best response to itself as mean invasion fitness cannot be further increased, being equal to 1 [thus eq. (2) implies eq. (1.B)].

The correspondence between uninvadability and Nash equilibrium allows us to lift two results from game theory to the problem of evaluating long-term evolutionary outcomes. To present these, let us focus on the case of no environmental fluctuations, when the uninvadable coalition is characterized by eq. (5). First, when is compact and the payoff function *ρ*(*x, p*) is continuous in *x* and *p*, then a Nash equilibrium *p*\* is guaranteed to exist (see Appendix A for a proof). Second, finding such a Nash equilibrium is computationally hard (e.g. Papadimitriou, 2007; Conitzer and Sandholm, 2008; Papadimitriou, 2014; Narahari, 2014, Theorem 2.1). Just finding the mixed Nash equilibrium in a two-player symmetric finite game belongs to the PPAD complexity class (e.g. Papadimitriou, 2007; Conitzer and Sandholm, 2008; Papadimitriou, 2014; Narahari, 2014, Theorem 2.1). There is strong evidence that problems in this class cannot be solved with a polynomial algorithm, i.e., PPAD≠P (Papadimitriou, 2007, p.39). While our population game does not necessarily reduce to a two-player symmetric game in mixed strategies (it can under certain cases), it is an infinite game and thus computational issues are likely to be harder. This does not mean that the uninvadable coalition can never be computed. If the invasion fitness has a special structure, it can be solved in certain cases. For instance, in succinct games (Papadimitriou, 2007), the analysis becomes more tractable, as seen in congestion games. In biological terms, this corresponds to models where competition for resources depends on some aggregate effect, such as foraging models with an ideal free distribution (e.g. Křivan et al., 2008).

The conclusion is twofold. First, one should be aware of the computational complexity of evaluating an uninvadable coalition and results from algorithmic game theory may be useful to evolutionary biology. Second, identifying conditions necessary for the emergence of adaptive polymorphism without computing the long term equilibrium is a valuable first-step to understanding polymorphism.

Box 2. Intensive versus extensive variables and explicit ecological models

When regulating factors are extensive variables they represent dynamical state variables that have reached some equilibrium (or more generally some attractor). For instance, when the regulating factors *e*_1_(*y*) and *e*_2_(*y*) in invasion fitness eq. (10) are extensive variables, they can be written to satisfy the following system of equations:

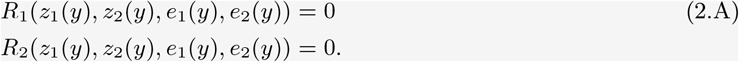

Here, *R*_1_ and *R*_2_ are mappings that characterise the equilibrium of the regulating variables, which are assumed to be locally asymptotically stable (see also Kisdi and Geritz, 2016, eq. 8). Since individuals interact with their environment via their production factor, the equilibrium values of both regulating factors *e*_1_(*y*) and *e*_2_(*y*) depend on both production factors *z*_1_(*y*) and *z*_2_(*y*). This could describe how competition for food resources *e*_1_(*y*) and breeding sites *e*_2_(*y*) is impacted by individuals’ efforts in foraging *z*_1_(*y*) or reproduction *z*_2_(*y*). An explicit example of an ecological map of the form eq. (2.A) and how extensive variables can be expressed in terms of intensive variables can be obtained as follows. Let us consider a population of iteroparous individuals that survive from one time step to the next with probability *z*_1_(*x*) and produce offspring according to a Beverton-Holt model of reproduction *z*_2_(*x*)*/*[1 + *γe*_1_(*y*) + *ηe*_2_(*y*)], where *z*_2_(*x*) is the baseline fecundity in the absence of density-dependence, *e*_1_(*y*) is the density of the focal species, *e*_2_(*y*) is the density of some other species, and *γ* and *η* are parameters tuning the importance of density-dependence. Invasion fitness of a mutant in the focal species then takes the form

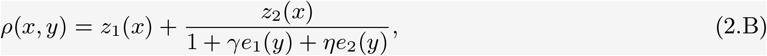

which is a Leslie-Gower model (Leslie and Gower, 1958, 1960). Since both *e*_1_(*y*) and *e*_2_(*y*) are extensive variables, we need maps describing population dynamics at equilibrium in these two species: *e*_1_(*y*) = *ρ*(*y, y*)*e*_1_(*y*) and *e*_2_(*y*) = *w*_2_(*y*)*e*_2_(*y*) where *w*_2_(*y*) is the fitness of an individual from the second species, which may depend on all quantities *z*_1_(*y*), *z*_2_(*y*), *e*_1_(*y*) and *e*_2_(*y*). Hence, in order to arrive at a system of the form of eq. (2.A), we can use

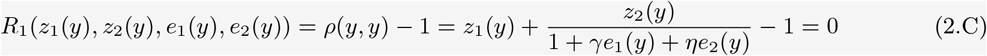

for the focal species, while for the second species we can use

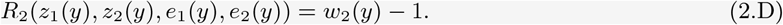

These allow us to solve for the population dynamic equilibrium of the focal species regardless of the functional form of eq. (2.D). Indeed, by solving eq. (2.C) for *e*_2_(*y*) and by substituting into eq. (2.B) produces

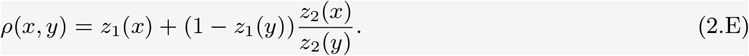

Thus, invasion fitness can be expressed in terms of mutant and resident production factors only, thus in terms of intensive variables. Interestingly, invasion fitness (2.E) can be seen as abiding to a lottery model for iteroparous reproduction with the following life-cycle (as in eq. 16 with *m* = 1): (a) a large fixed number of adults produce a large number of juveniles with fecundity *z*_2_ and then each adult either survives or dies independently (with probabilities *z*_1_ and 1 − *z*_1_, respectively); (b) density-dependent competition occurs among juveniles for the breeding spots vacated by the death of adults so that the population is regulated back to the fixed adult size. Such life-cycles are standard in population genetics. Assuming further *z*_1_(*x*) = 0 gives the Wright-Fisher model (Ewens, 2004; Hartl and Clark, 2007). These connections illustrate that standard models of population genetics can implicitly incorporate ecological feedbacks, even when these are not made explicit (as is well established, e.g. Crow and Kimura, 1970; Nagylaki, 1992). In such models, ecological feedbacks do not modify the selection pressures being studied, which is why they can be omitted without loss of generality.

Box 3. Directional selection and NTDS in class structured populations

We consider here heterogeneous populations where individuals can belong to a finite number of classes, where 𝒞 denotes the set of possible classes (e.g., genetic background, condition, age, environmental state). Assuming that this heterogeneity occurs either only within-generations (e.g. stage) or only between-generations–in which case heterogeneity is independently and identically distributed (e.g. temporal fluctuations in climate)–invasion fitness is given by

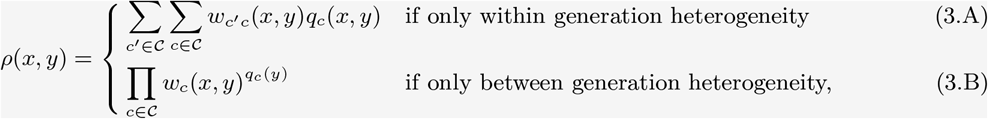

where mutant individuals express trait value *x* while the resident population has trait *y*. In eq. (3.A), *w*_*c*_′_*c*_(*x, y*) is the expected number of offspring of class *c*^′^ (including the surviving self) produced by a mutant in class *c*, and *q*_*c*_(*x, y*) is the probability that a mutant is in class *c* (see Appendix G.1 or Lehmann et al., 2016. We assume that the matrix with elements *w*_*c*_′_*c*_(*x, y*) is primitive and the branching process is non-singular (each individual does not produce exactly one successful offspring), which ensures that the invasion dichotomy holds (e.g. Harris, 1963, p. 38-41). In eq. (3.B), *w*_*c*_(*x, y*) is the expected number of offspring produced by a mutant in class or environmental state *c* and *q*_*c*_(*y*) is the probability of occurrence of environmental state *c* ∈ 𝒞, which depends only on resident traits (as in standard invasion analysis, e.g., Ferrière and Gatto, 1995). Note that owing to between-generation heterogeneity, there is only one type of offspring so *w*_*c*_(*x, y*) depends only on the environmental state *c* in which the individual reproduces. We show in Appendix G.3 how these class-specific quantities can be expressed in terms of regulation and production factors such that invasion fitness in both cases of heterogeneity can be conceptualised as a mapping from fitness factors to geometric growth ratio (i.e. as in eq. 9).

Using eqs. (3.A)-(3.B), the selection gradient can be expressed as

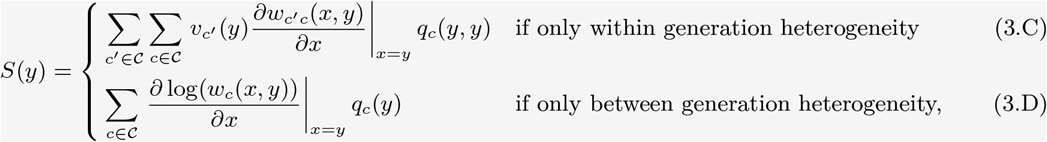

where *v*_*c*_(*y*) is the reproductive value of an individual in class *c* in a (monomorphic) resident population *y*, satisfying *v*_*c*_(*y*) = ∑_*c*_′_∈𝒞_*v*_*c*_′ (*y*)*w*_*c*_′_*c*_(*y, y*) while *q*_*c*_′ (*y, y*) = ∑_*c* ∈𝒞_*w*_*c*_′_*c*_(*y, y*)*q*_*c*_(*y, y*) (eq. 3.C is a standard result, e.g., Taylor and Frank, 1996; Rousset, 2004; Lehmann et al., 2016 and eq. 3.D is derived in Appendix G.2). The singular trait value *y*\* thus reflects a trade-off of fitness effects among different classes, which are appropriately weighted according to how often an individual is in class *c* (via *q*_*c*_(*y, y*) or *q*_*c*_(*y*)) and offspring reproductive value under within generation heterogeneity (via *v*_*c*_′ (*y*) in eq. 3.C).

The interaction coefficient, meanwhile, can be written as,

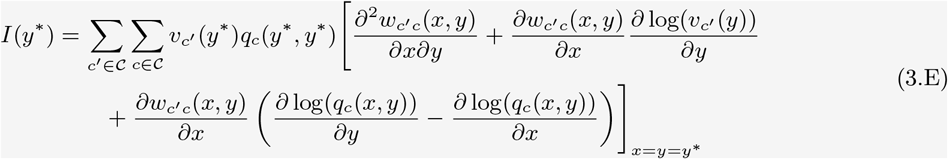

for within generation heterogeneity, and

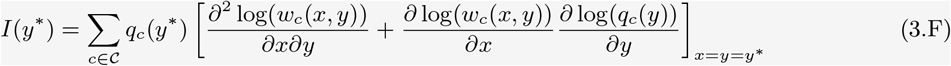

for between generation heterogeneity (Appendix G for derivation). The first terms between square brackets in eqs. (3.E) and (3.F) show that NTDS can simply be due to negative interaction effects on class specific fitness components (i.e. via *∂*^2^*w*_*c*_′_*c*_(*x, y*)*/*(*∂x∂y*) and *∂*^2^ log(*w*_*c*_(*x, y*))*/*(*∂x∂y*)). The rest of eqs. (3.E) and (3.F) highlight how population heterogeneity opens new pathways for NTDS. These can be understood by assuming that mutant trait expression increases mutant fitness in a given class (*∂w*_*c*_′_*c*_(*x, y*)*/∂x >* 0 or *∂* log(*w*_*c*_(*x, y*)*/∂x >* 0). A joint increase in the resident trait value then tends to have negative effects on mutant invasion fitness when such an increase reduces: (i) the reproductive value of offspring produced in that class under within generation heterogeneity (via *∂* log(*v*_*c*_′ (*y*))*∂y <* 0 in eq. 3.E) and/or (ii) the probability that mutants belong to the class in which their fitness is enhanced in eq. 3.F, and where this effect is measured relative to the mutant’s effect on this probability in eq. 3.E. This comes from the fact that in populations with within generation heterogeneity, natural selection depends on class specific fitness differences (section Appendix G.1 for more details).

## Footnotes

1 In invasion analyses of evolutionary ecology, invasion fitness is often written as *ρ*(*x, µ*(𝒫 (*µ*_p_))), thus emphasizing that invasion fitness depends on the full environment *µ* of the resident population, which itself is determined by the set of segregating morph 𝒫 (*µ*_p_) (e.g. Geritz et al., 1998; Metz et al., 2008; Metz, 2011). But since this set of morphs is determined by *µ*_p_, we can write invasion fitness as *ρ*(*x, µ*_p_) as a shortcut, keeping in mind that the full environment *µ* would need to be characterized to compute invasion fitness (section 3.2 and Box 2 make these points more explicit for specific models).

2 Strictly speaking this increase is due to the interaction between mutant and resident trait values only; for brevity we omit to systematically refer to “interaction” in the paper.

3 If survival is incomplete during dispersal so that there is a cost to dispersal, *m* should be interpreted as the backward migration rate (Nagylaki, 1992), i.e., *m* = *ds*_d_*/*[(1 *− d*) + *ds*_d_], where *d* is the probability of dispersal out of the natal site and *s*_d_ the probability of survival during dispersal. All forthcoming models and results of the paper carry through with *m* taken as the backward migration probability, but for simplicity of presentation we simply refer to *m* as the migration probability.

4 Denote by *N* (*t*) the amount of resource on a fast time scale *t* and postulate the dynamics 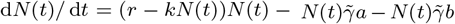 for some intrinsic resource growth rate *r*, strength of competition *k*, and extraction rate 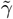. At equilibrium, the amount of resource is 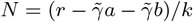, which by setting *α* = *r/k* and 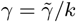 becomes *N* = (*α − γa − γb*), as required.

